# Lgr5-mediated restraint of β-catenin is essential for B-lymphopoiesis and leukemia-initiation

**DOI:** 10.1101/2020.03.12.989277

**Authors:** Kadriye Nehir Cosgun, Mark E. Robinson, Gauri Deb, Xin Yang, Gang Xiao, Teresa Sadras, Jaewoong Lee, Lai N. Chan, Kohei Kume, Maurizio Mangolini, Janet Winchester, Zhengshan Chen, Lu Yang, Huimin Geng, Shai Izraeli, Joo Song, Wing-Chung Chan, Andrew G. Polson, Hassan Jumaa, Hans Clevers, Markus Müschen

**Author notes:** **For correspondence:** Markus Müschen, MD-PhD, Department of Systems Biology, City of Hope Comprehensive Cancer Center, 1218 South Fifth Ave, Monrovia, CA 91016.

## Abstract

Upon productive immunoglobulin gene rearrangement, expression of a functional pre-B cell receptor (pre-BCR) initiates positive selection of pre-B cells, clonal expansion and self-renewal^1-2^. Studying mechanisms driving this first wave of B-lymphopoiesis, we identified the G-protein coupled receptor Lgr5 as an essential initiator of positive selection. Lgr5 was extensively studied as determinant of stem cell populations in multiple tissues^3-6^, but not in B-cells. While undetectable throughout the hematopoietic system, positively selected pre-B cells were marked with a sharp peak of Lgr5 expression. Conditional deletion of *Lgr5* preceding the pre-BCR checkpoint induced negative selection and complete abortion of B-cell development. Proteomic studies of *Lgr5*-ablation revealed massive (>250-fold) accumulation of β-catenin and suppression of MYC. *Lgr5*-deficient pre-B cells fully recovered by concurrent β-catenin-deletion, demonstrating a central role of Lgr5-mediated restraint of β-catenin at the pre-BCR checkpoint. In other cell types, β-catenin/TCF4 complexes drive transcriptional activation of MYC^7-9^. Instead of TCF4, proximity-based interactome studies in pre-B cells identified the B-lymphoid transcription factors IKZF1 and IKZF3^10-11^ as β-catenin-binding partners, which had the opposite effect and caused transcriptional repression of *MYC*. On positively selected pre-B cells, Lgr5 prevented accumulation of β-catenin and formation of complexes with IKZF1 and IKZF3, which relieved transcriptional repression of MYC. Activating β-catenin-mutations are common throughout all main types of cancer^7-8^, but were conspicuously absent in pre-B leukemia (B-ALL). Like pre-B cells, B-ALL cells were uniquely sensitive to genetic and pharmacological β-catenin hyperactivation, which recapitulated the effects of *Lgr5*-deletion and compromised colony formation and leukemia-initiation. A new LGR5 antibody-drug conjugate targeted leukemia-initiating cells in patient-derived B-ALL and achieved long-term disease-control. Likewise, small molecule hyperactivation of β-catenin selectively killed B-ALL but not other cell types. Hence, Lgr5-mediated restraint of β-catenin activation is essential for B-lymphopoiesis and revealed an unexpected vulnerability that can be leveraged for the treatment of drug-resistant B-ALL.

## Lgr5 is essential for the initiation of B-lymphopoiesis

Once early B-lymphocyte precursors have productively rearranged immunoglobulin (Ig) V, D and J gene segments, expression of a functional Ig μ-heavy chain as part of the pre-B cell receptor (pre-BCR) results in a strong positive selection signal to initiate clonal expansion and the first wave of B-lymphopoiesis^1, 2^. To study mechanisms of positive selection and pre-B cell self-renewal, we analyzed gene expression changes that are induced at the onset of Ig μ-heavy chain signaling. Among the most prominent changes at the pre-BCR checkpoint, we identified upregulation of the Leucine-rich repeat-containing G-protein coupled receptor 5 (Lgr5; **Fig. 1a** and **b, Extended data figure 1a**). While mRNA levels of Lgr5 were low or undetectable throughout the entire spectrum of hematopoietic lineages, the pre-BCR checkpoint was marked by a sharp peak of Lgr5 expression (Hardy Fractions C-D; **Fig. 1c, Extended data figure 1b-c**). Lgr5 was extensively studied as determinant of adult stem cell populations in multiple tissues including intestinal^3^, kidney, liver^4^ and gastric tissues as well as ear and hair follicles^*5*^. For instance, in the colon crypt, Lgr5 marks quiescent stem cell populations that promote tissue regeneration and give rise to the main lineages of the colon epithelium^*6*^. Mechanistically, Lgr5 is thought to interact with Rspo1 to potentiate WNT signaling by stabilizing Frizzled (FZD) surface receptors of WNT^12^, induce assembly of disheveled (DVL2)^13-16^ and β-arrestin (ARRB2)^17-20^ scaffolds, which ultimately results in accumulation of β-catenin.

**Figure 1:**
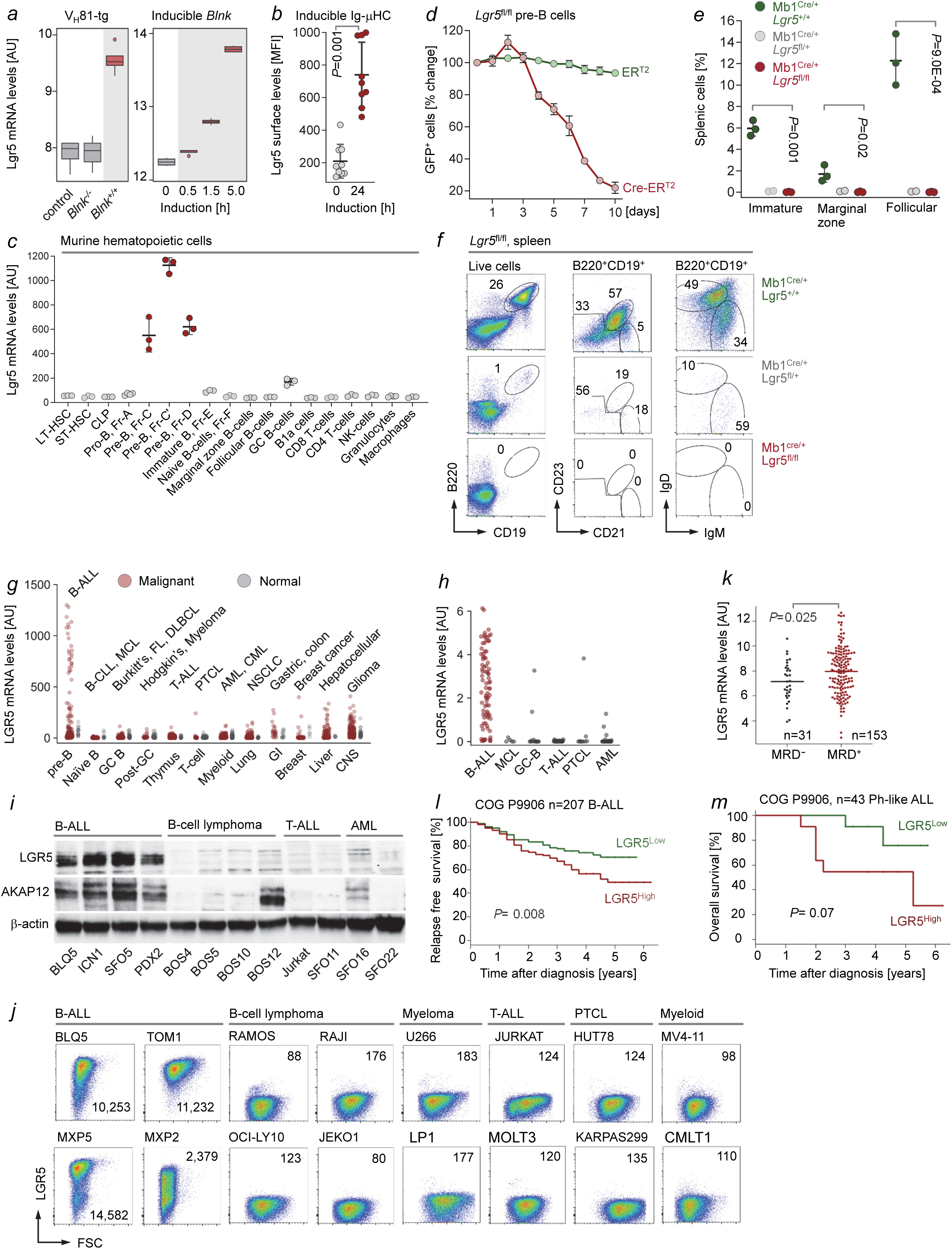
Identification of Lgr5 as an essential requirement for B-lymphopoiesis. (**a**) Microarray analyses (GSE53896) of Lgr5 mRNA levels in FACS-sorted B220^+^ CD19^+^ pre-B cell fractions from VH81x transgenic mice with functional (*Blnk*^+/+^, n=4) or non-functional (*Blnk*^-/-^, n=4) pre-BCR signaling chains are shown (left). Non-VH81x transgenic *Rag1*^-/-^ pro-B cells were used as controls (n=3). RNA-seq analysis of *Blnk*^-/-^ pre-B-cells that were reconstituted with 4-hydroxy-tamoxifen (4-OHT) inducible *Blnk* for inducible activation of pre-BCR signaling (right). Lgr5 mRNA levels at 0 to 5 hours after 4-OHT addition (GSE73562) were indicated. (**b**) *Rag2*^-/-^ pro-B cells were transduced with a 4-OHT-inducible Ig μ-heavy chain (μHC) to initiate pre-BCR signaling. Lgr5 surface levels were measured by flow cytometry before and 24 hours after induction (n=9). (**c**) ImmGen microarray data sets (GSE15907, GSE37448) were analyzed for Lgr5 mRNA levels in 18 murine hematopoietic populations including 10 B-cell subsets. Pre-B cell stages (Fractions C-D) are highlighted in red. (**d**) IL7-dependent pre-B cells from the bone marrow of *Lgr5*^fl/fl^ mice were transduced with GFP-tagged, 4-OHT-inducible Cre (Cre-ER^T2^) or empty vector controls (ER^T2^). Following 4-OHT induction, percentages of GFP^+^ cells, normalized to fraction of GFP^+^ cells before induction, were monitored by flow cytometry (n=3). Immature B cells (CD21^-^ CD23^-^), marginal zone B cells (CD21^+^ CD23^-^) and follicular B cells (CD21^+^ CD23^+^) were analyzed by flow cytometry of splenic tissue from *Mb1*^Cre/+^ *Lgr5*^+/+^, *Mb1*^Cre/+^ *Lgr5*^fl/+^, *Mb1*^Cre/+^ Lgr5^fl/fl^ mice (n=3). (**e**) Relative frequencies and (**f**) representative FACS stainings are shown. Microarray data for LGR5 mRNA levels were studied in hematological malignancies and solid tumors. In (**g**) LGR5 mRNA levels are shown for 7 hematological malignancies and 5 solid tumor types (red dots) as well as their normal counterpart or cell of origin (gray dots)^68^. In (**h**), LGR5 mRNA levels in B-ALL samples (red) were compared to other hematological malignancies (gray)^69^. (**i**) Western blot analyses were performed to measure protein levels of LGR5 and AKAP12 in a panel of patient derived xenografts (PDX) and Jurkat T-ALL cells. (**j**) Surface levels of LGR5 expression was determined by flow cytometry in a panel human B-ALL, B-cell lymphoma, multiple myeloma, T-ALL, peripheral T-cell lymphoma (PTCL) and myeloid leukemia cell lines and PDX samples. (**k**) Minimal residual disease (MRD) status from patients with pediatric high-risk B-ALL (COG P9906) was determined by flow cytometry on day 29 post treatment. LGR5 expression levels were compared in patients with MRD^+^ versus MRD^-^ status (*P*=0.025). (**l**) B-ALL patients from the same clinical cohort were segregated into two groups based on whether LGR5 mRNA levels were higher (LGR5^high^) or lower (LGR5^low^) than median expression. Relapse-free survival (RFS) was assessed in the two groups by Kaplan-Meier analysis (log-rank test, *P*=0.008). (**m**) This cohort included a subset of *Ph*-like ALL (n=43) patients. Overall survival (OS) was assessed in the LGR5^high^ and LGR5^low^ groups by Kaplan-Meier analysis, however the difference in OS did not reach statistical significance (log-rank test, *P*=0.07).

A role of Lgr5 was not previously examined in B-lymphocyte development. To study the functional significance of sharp upregulation of Lgr5 at the pre-BCR checkpoint (Hardy Fractions C-D), we tested the consequences of inducible Cre-mediated deletion of *Lgr5* in developing pre-B cells. Under cell culture conditions, inducible *Lgr5*-deletion caused rapid loss of competitive fitness of pre-B cells (**Fig. 1d**). In a conditional mouse model for deletion of *Lgr5* from earliest stages of B-cell development (*Mb1*-Cre), pro-B cells (Hardy Fractions A-B) developed normally. However, B cell precursors could not progress past the pre-BCR checkpoint resulting in complete abortion of B-cell development in these mice (**Fig. 1e-f, Extended data figure 2**). Even deletion of only one allele of *Lgr5* was sufficient to cause near-complete ablation of B-cell development, suggesting that even moderate reduction of *Lgr5* gene expression may compromise positive selection at the pre-BCR checkpoint (**Fig. 1e-f, Extended data figure 2**). Interestingly, while deletion of *Lgr5* compromised B-lymphopoiesis *in vivo* and competitive fitness *in vitro*, loss of Lgr5 did not significantly increase apoptosis and cell death (**Extended data figure 9d, f**). In addition, when deletion of *Lgr5* was induced in mature B-cells after the pre-BCR checkpoint (*Cd21*-Cre; **Extended data figure 3**) or in germinal center B-cells (*Aicda*-Cre; **Extended data figure 4**), loss of *Lgr5* only caused a modest reduction of marginal zone B-cells. All other mature B-cell subsets, including B1 cells remained unaffected. Lgr5-deficient germinal center (GC) B-cells underwent normal affinity maturation and were able to develop antigen-specific B cells in response to immunization (**Extended data figure 4**). Consistent with a sharp peak of gene expression in pre-B cells (Hardy Fractions C-D), these findings indicate that Lgr5 is uniquely required for positive selection at the pre-BCR checkpoint. Once mature B-cell populations have formed, they no longer depend on Lgr5 function for maintenance and affinity maturation in humoral immune responses.

**Figure 2:**
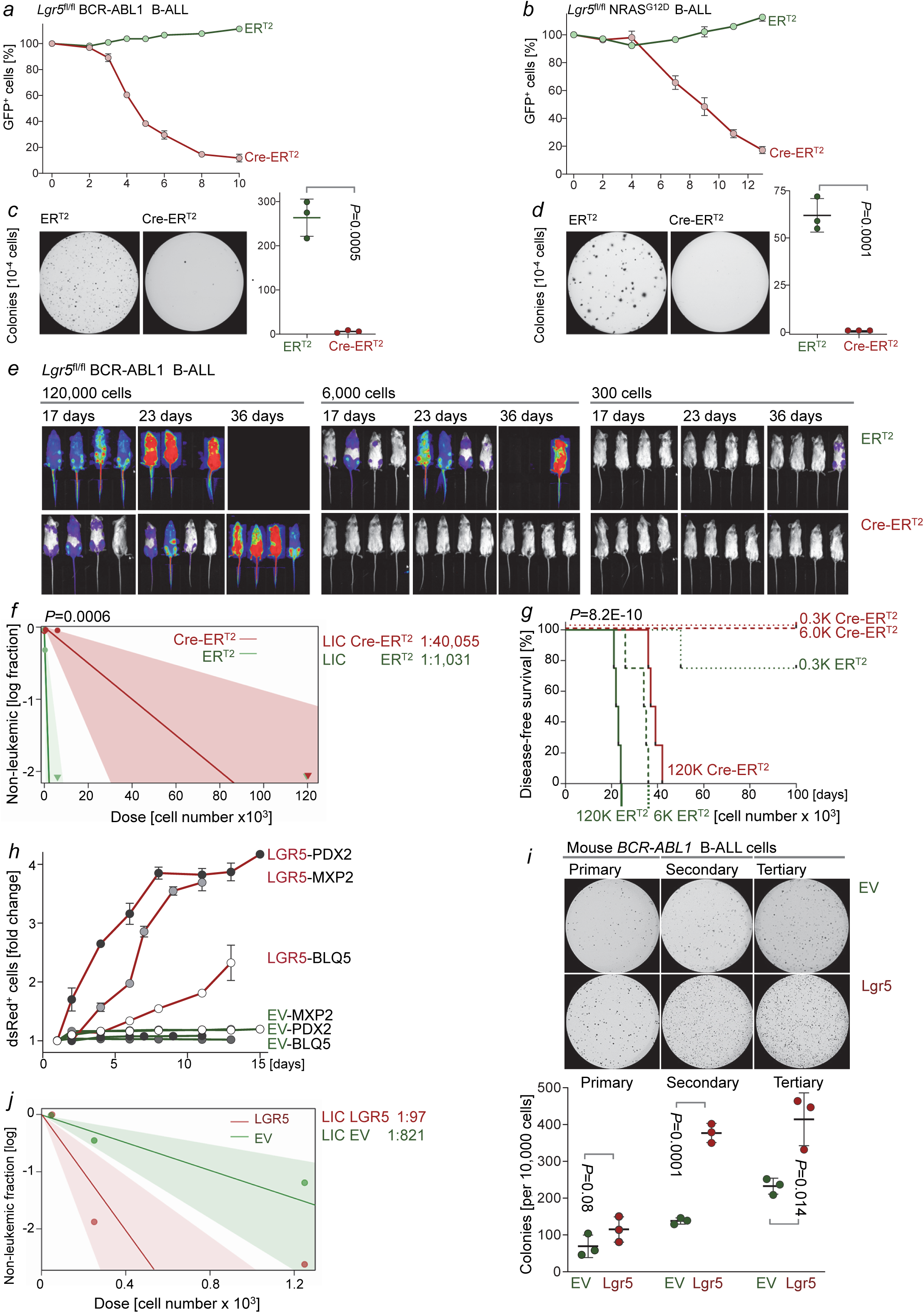
Lgr5 is essential for B-ALL leukemia-initiation. Lgr5^fl/fl^ pre-B cells were retrovirally transformed by (**a**) BCR-ABL1 or (**b**) NRAS^G12D^ and B-ALL cells were transduced with GFP-tagged, 4-OHT-inducible Cre (Cre-ER^T2^) or empty vector (ER^T2^) control. Following 4-OHT induction, frequencies of GFP^+^ cells were monitored by flow cytometry (n=3), normalized to frequencies measured on day 0. (**c**) BCR-ABL1 or (**d**) NRAS^G12D^ *Lgr5*^fl/fl^ B-ALL cells carrying ER^T2^ or Cre-ER^T2^ were plated on semisolid methylcellulose agar 2 days after 4-OHT induction. Colonies were imaged and counted after 10 days (n=3). (**e**) BCR-ABL1-driven *Lgr5*^fl/fl^ B-ALL cells carrying ER^T2^ or Cre-ER^T2^ and firefly luciferase were transplanted into sub-lethally irradiated (2 Gy) NSG recipients at three dose levels (300 cells, 6,000 cells, 120,000 cells; n=4). Leukemia burden and disease progression were measured by bioluminescence imaging. (**f**) Extreme limiting dilution analysis (ELDA) was performed to calculate leukemia-initiation capacity. LIC-frequencies of 1 in 1,031 cells were measured in ERT2-expressing cells compared to 1 in 40,055 cells in Cre-ERT2 expressing cells (*P*=0.0006). (**g**) Kaplan-Meier analysis shows overall survival in each group and dose level (n=4; *P*=8.2E-10; log-rank test). (**h**) Patient-derived B-ALL xenografts (PDX2, MXP2, BLQ5) were transduced with ds-Red-tagged LGR5 or EV control vectors. Competitive advantage or depletion of ds-Red^+^ cells were monitored by flow cytometry, normalized to the percentage of ds-Red^+^ cells at day 0. (**i**) BCR-ABL1 transformed mouse B-ALL cells were transduced with doxycycline-inducible LGR5 or EV control. 10,000 cells were plated on semisolid methylcellulose agar and doxycycline was added. Primary colonies were counted and imaged after 10 days and replated in serial replating assays (n=3). (**j**) Flow-sorted ds-Red^+^ PDX2 B-ALL cells carrying LGR5-dsred or EV-dsRed were transplanted into sub-lethally irradiated (2 Gy) NSG recipients (n=7). ELDA analysis was performed to calculate LIC frequencies, 1 in 821 in the EV group, compared to 1 in 97 in LGR5-overexpressing B-ALL cells (*P*=0.0098).

**Figure 3:**
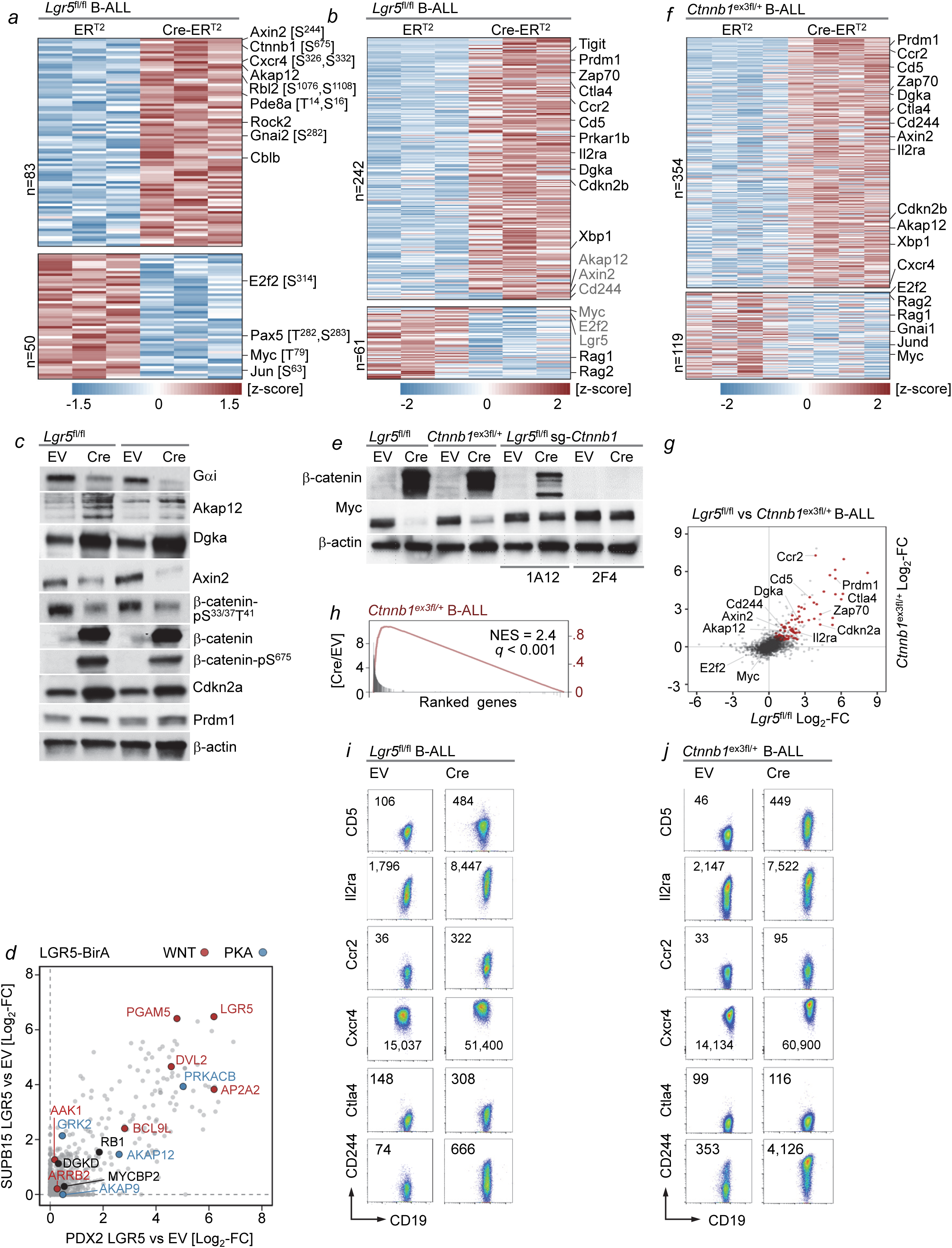
LGR5 functions as an essential negative feedback regulator of β-catenin in B-ALL cells. (**a**) *Lgr5*^fl/fl^ BCR-ABL1 B-ALL cells carrying 4-OHT-inducible Cre-ER^T2^ or ER^T2^ empty vector control were treated with 4-OHT for 3 days. Differentially phosphorylated proteins were identified by mass spectrometry (n=3). (**b**) RNA-seq analysis of *Lgr5*^fl/fl^ BCR-ABL1 B-ALL cells carrying Cre-ER^T2^ or ER^T2^ were treated with 4-OHT for 3 days and gene expression changes were plotted as heatmap. (**c**) Western blot analysis for *Lgr5*^fl/fl^ BCR-ABL1 B-ALL cells carrying Cre-ER^T2^ or ER^T2^ was performed to validate gene expression changes at the protein level. Western blots were performed Gαi, Akap12, Dgkα, Axin2, (destabilized) β-catenin-pS^33^/S^37^/T^41^, (activated) β-catenin-pS^675^, global β-catenin, Cdkn2a and Prdm1, using β-actin as loading control. (**d**) To identify LGR5-interaction partners, we generated C-terminal fusion proteins between the cytoplasmic tail of LGR5 and the bacterial biotin-ligase BirA. Proximity-based analyses of proteins that were biotinylated by LGR5-BirA fusions (Bio-ID) identified dishevelled2 (DVL2), its scaffold β-arrestin (ARRB2), as well as the DVL2 clathrin adapter AP2 (AAK1, AP2A2, AP2B1) as central interaction partners. Bio-ID analysis were performed for human PDX2 (x-axis) and SUPB15 (y-axis) B-ALL cells. WNT/β-catenin-related proteins are labeled in red, PKA-related proteins in blue. (**e**) *Lgr5*^fl/fl^ and *Ctnnb1*^ex3fl/+^ B-ALL cells and *Lgr5*^fl/fl^ B-ALL cells carrying retroviral Cas9 for deletion of *Ctnnb1* (*Lgr5*^fl/fl^ sg-*Ctnnb1*) were transduced with Cre (for deletion of *Lgr5* or stabilization of β-catenin) or EV. Western blot analysis was performed to measure β-catenin and Myc protein levels, using β-actin as loading control. For Cas9-mediated deletion of *Ctnnb1*, results for two sg-*Ctnnb1* guides, 1A12 and 2F4, are shown. (**f**) RNA-seq analysis of *Ctnnb1*^ex3fl/+^ BCR-ABL1 B-ALL cells carrying ER^T2^ or Cre-ER^T2^ was performed 1 day after 4-OHT addition, gene expression changes shown as heatmap. Gene expression changes [log2-fold] in *Lgr5*^fl/fl^ (x-axis) and *Ctnnb1*^ex3fl/+^ (y-axis) B-ALL cells were compared in a scatter plot (**g**) and through GSEA analysis (**h**). Common gene expression changes in *Lgr5*^fl/fl^ (**i**) and *Ctnnb1*^ex3fl/+^ (**j**) B-ALL cells that affected cell surface receptors were validated by flow cytometry. FACS dot plots for double stainings for Cd19 with Cd5, Il2ra (Cd25), Ccr2, Cxcr4, Ctla4 and Cd244 (2B4) are shown without (EV) or 3 days after (Cre) 4-OHT-mediated induction of Cre-ER^T2^. Numbers in FACS plots denote mean fluorescence intensities.

**Figure 4:**
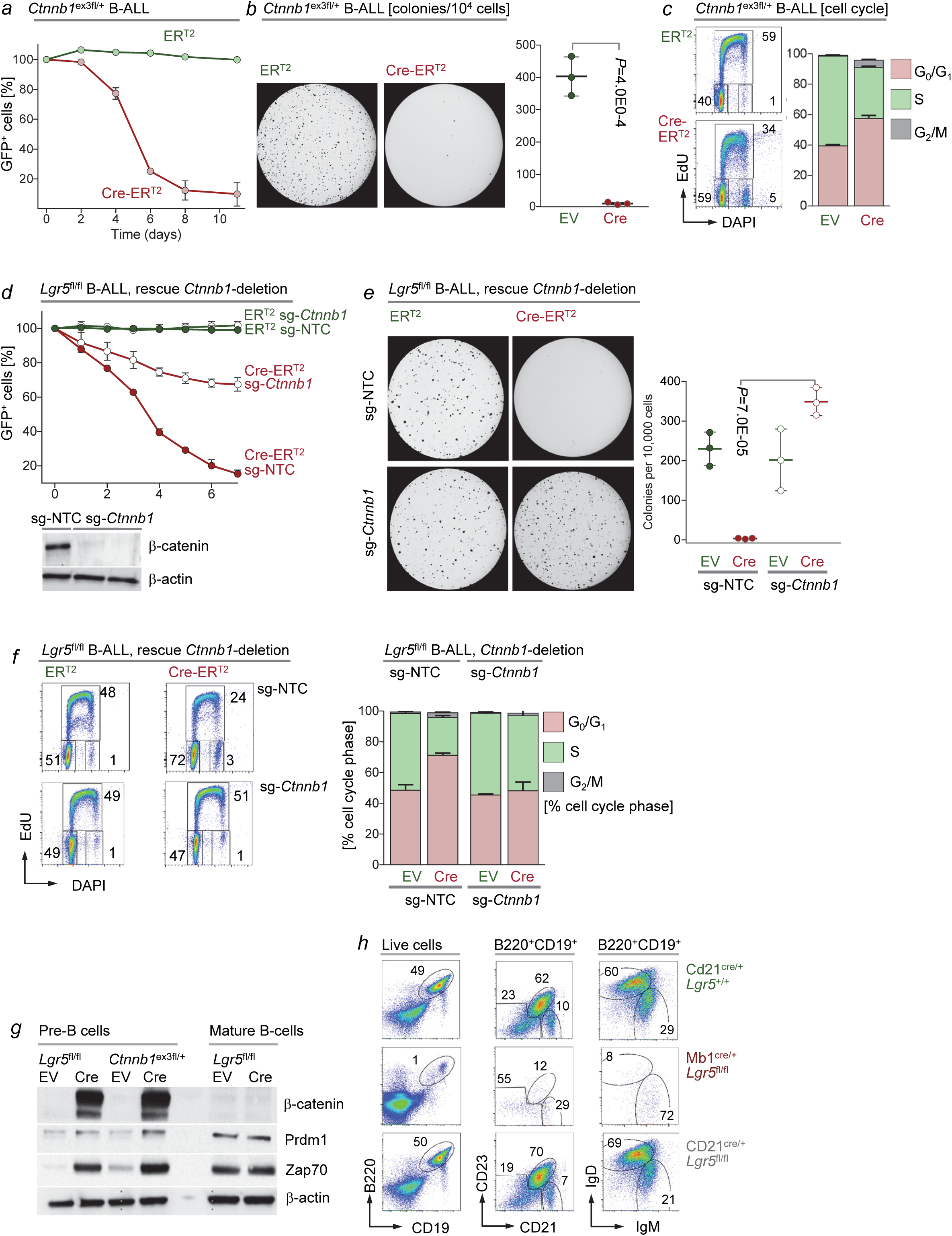
Lgr5-mediated negative regulation of β-catenin is essential for B-lymphopoiesis and B-ALL initiation. (**a**) Bone marrow pre-B cells from *Ctnnb1*^ex3fl/+^ mice were transformed by BCR-ABL1 and transduced with GFP-tagged Cre-ER^T2^ or ER^T2^ constructs. Changes of percentages of GFP^+^ cells were monitored for 12 days upon 4-OHT addition (n=3). (**b**) 10,000 *Ctnnb1*^ex3fl/+^ BCR-ABL1 B-ALL cells carrying Cre-ER^T2^ or ER^T2^ were plated for colony formation assays 2 days after 4-OHT treatment. Representative images and colony numbers are shown at 10 days after plating (n=3). (**c**) Cell cycle phases of *Ctnnb1*^ex3fl/+^ BCR-ABL1 B-ALL cells carrying Cre-ER^T2^ or ER^T2^ were measured by EdU incorporation in combination with DAPI staining. (**d**) *Lgr5*^fl/fl^ BCR-ABL1 B-ALL cells were transduced with GFP-tagged Cre-ER^T2^ or ER^T2^ and transduced with Cas9 and guides for non-targeting controls (sg-NTC) or guides targeting *Ctnnb1* (sg-*Ctnnb1*). Deletion of *Ctnnb1* was confirmed by Western blot (**d**) in clonal cell lines that grew out from single clones (**Extended data figure 10**). Competitive fitness of *Lgr5*^fl/fl^ B-ALL clones was measured based on deletion of *Lgr5* (Cre-ER^T2^) and/or deletion of *Ctnnb1* (sg-*Ctnnb1*). (**e**) *Lgr5*^fl/fl^ B-ALL cells with (Cre-ER^T2^) or without (ER^T2^) deletion of *Lgr5* in combination with (sg-*Ctnnb1*) or without (sg-NTC) deletion of *Ctnnb1* were plated for colony forming assays at 10,000 cells per plate. Colonies were imaged and counted 10 days after plating (n=3). (**f**) Cell cycles phases in *Lgr5*^fl/fl^ B-ALL cells with (Cre-ER^T2^) or without (ER^T2^) deletion of *Lgr5* in combination with (sg-*Ctnnb1*) or without (sg-NTC) deletion of *Ctnnb1* were studied by Edu incorporation and DAPI staining 3 days after 4-OHT treatment for Cre-mediated deletion of *Lgr5*. (**g**) Western blot analyses of β-catenin, Prdm1, Zap70 protein levels in bone marrow pre-B cells from *Lgr5*^fl/fl^ or *Ctnnb1*^ex3fl/+^ mice in comparison to mature splenic B-cells from *Lgr5*^fl/fl^ mice without (EV) and after activation of Cre. (**h**) Representative FACS staining of mature B cell populations in the spleens of *Cd21*^Cre/+^ Lgr5^+/+^ mice (no deletion of *Lgr5*), *Mb1*^Cre/+^ *Lgr5*^fl/fl^ mice (early deletion of *Lgr5* prior to pre-BCR checkpoint) and *Cd21*^Cre/+^ *Lgr5*^fl/fl^ mice (late deletion of *Lgr5*, past the pre-BCR checkpoint) are shown.

## Lgr5 expression predicts poor clinical outcomes in patients with B-ALL

While Lgr5 contributes to self-renewal of normal epithelial stem cells and tissue regeneration, this is also the case for cancer-initiating cells. For instance, lineage-tracing experiments revealed Lgr5 as determinant of tumor-initiation in intestinal adenomas^21^. In addition, Lgr5 marks colon cancer-initiating cells^22^, promotes metastasis^23^ and represents a therapeutic target for eradication of colon cancer stem cells (NCT02726334, NCT03526835)^24,25^. For this reason, we studied LGR5 mRNA levels in patient-derived samples of B-cell acute lymphoblastic leukemia (B-ALL), the tumor that originates from pre-B cells, compared to common solid tumors and hematopoietic malignancies. While LGR5 is expressed in multiple solid tumors, including colon cancer, gastric, hepatocellular, breast cancer and glioma, we found the highest LGR5 mRNA levels in B-ALL samples (**Fig. 1g**). Consistent with confinement of LGR5 expression to the pre-B cell stage, LGR5 mRNA levels were high in most cases of B-ALL but low or undetectable in mature B-cell malignancies derived from naïve B-cells (chronic lymphocytic leukemia, mantle cell lymphoma), GC B-cells (Burkitt’s lymphoma, follicular lymphoma, diffuse large B-cell lymphoma) as well as post-GC B-cell malignancies (Hodgkin’s disease, multiple myeloma; **Fig. 1g-h**). Comparing LGR5 mRNA levels in cytogenetic B-ALL subtypes, we found high levels throughout all B-ALL subtypes, while *MLL*-rearranged B-ALL, mostly originating from pro-B cells, expressed LGR5 at lower levels (**Extended data figure 1d**). High expression levels of LGR5 in patient-derived B-ALL cells were confirmed at the protein level by Western blot and flow cytometry (**Fig. 1i-j, Extended data figure 1e**). To determine whether LGR5 expression may predict clinical outcomes in patients with lymphoid malignancies, LGR5 mRNA levels were measured at the time of diagnosis in one clinical trial for patients with B-ALL (P9906, n=207; **Fig. 1k-m**), six clinical cohorts of patients with mature B-cell malignancies and one T-cell lymphoma trial (**Extended data figure 5**). In each clinical cohort, patients were then segregated into two groups based on higher or lower than median LGR5 mRNA levels. Comparing persistence of minimal residual disease (MRD), overall survival and relapse-free survival between the two groups, higher LGR5 mRNA levels predicted poor clinical outcome in B-ALL (**Fig. 1k-m**) but not in any other hematological malignancies (**Extended data figure 5**). Mirroring Lgr5-functions during early B-cell development, these results suggest a disease-specific role of LGR5 in B-ALL.

**Figure 5:**
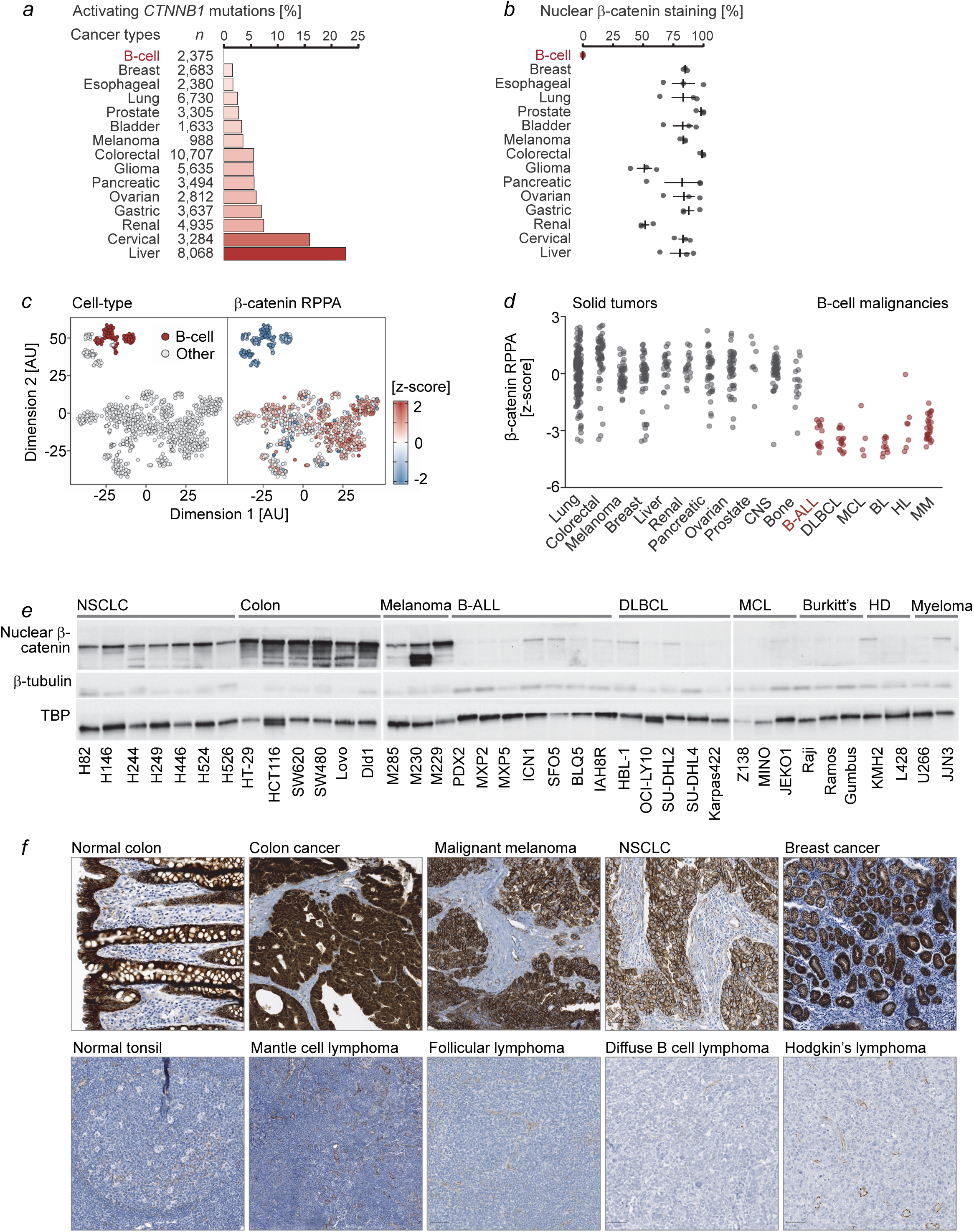
B-lymphoid malignancies are exempt from oncogenic activation of β-catenin signaling. (**a**) The frequencies of activating mutations of β-catenin (hot spot mutations in β-catenin exon 3: D^32^, S^33^, G^34^, S^37^, T^41^, S^45^) are depicted for 15 types of cancer including B-cell malignancies and solid tumors. Analyzing data from cancer 66,820 samples (COSMIC, *49*), we identified 4,971 activating *CTNNB1* mutations (on average 7.4%) but none in 2,375 B-cell malignancies. (**b**) 519 tumor samples representing 17 types of cancer were stained for β-catenin expression using three different antibodies (1,521 experiments total) and analyzed by immunohistochemistry. Among 1,488 solid tumor samples, in 1,202 cases (81%) β-catenin expression was detected, compared to none in 33 B-cell lymphoma samples. (**c**) Global transcriptional profiles (RNA-seq) segregated the 110 B-cell tumor cell lines from the 847 cancer cell lines (tSNE plots; left), superimposing cell-line matched reverse-phase protein array (RPPA) data for β-catenin onto these clusters revealed strikingly low β-catenin levels (heatmap for β-catenin protein levels, right). (**d**) β-catenin levels measured by RPPA were plotted for cancer cell lines representing 11 solid tumors (gray) and 6 B-cell malignancies (red). (**e**) Western blot analyses on nuclear fractions of non-small lung carcinoma (NSCLC), colon cancer, malignant melanoma leukemia and B-ALL, diffuse large B-cell lymphoma (DLBCL), mantle cell lymphoma (MCL), Burkitt’s, Hodgkin’s disease (HD) and multiple myeloma cell lines for β-catenin, β-tubulin and TBP. Western blots of the cytoplasmic fractions from the same cell lysates are shown in Extended data figure 11a. (**f**) Immunohistochemical staining for β-catenin was performed on normal colon (n=2) and tonsil (lymphoid follicle, n=1), as well as colon cancer (n=25), malignant melanoma (n=5), NSCLC (n=15) and breast cancer in comparison to mantle cell lymphoma (n=26), follicular lymphoma (n=38), DLBCL (n=35) and Hodgkin’s (n=44) disease.

## Lgr5 is essential for B-ALL leukemia-initiation

To model common subtypes of B-ALL^26^, we transduced pre-B cells from *Lgr5*^fl/fl^ mice^12^ with BCR-ABL1 or NRAS^G12D^ oncogenes and tamoxifen-inducible Cre (Cre-ER^T2^) or EV (ER^T2^; **Fig. 2**). Induction of Cre-mediated deletion of *Lgr5* rapidly reduced competitive fitness of B-ALL cells under cell culture conditions (**Fig. 2a-b, Extended data figure 6a-b**) and abolished the ability of B-ALL cells to form colonies (**Fig. 2c-d**). When saturating numbers of full-blown B-ALL cells were injected into sublethally irradiated NSG mice, Cre-mediated deletion of *Lgr5* substantially prolonged survival of transplant recipient mice (**Extended data figure 6c-d**). However, ultimately all mice died from fatal leukemia, suggesting that Lgr5 is not required for maintenance of established leukemia. In a serial transplant experiment based on limiting dilution of cell numbers (300, 6,000 and 120,000 injected B-ALL cells), Cre-mediated deletion of *Lgr5* resulted in a profound defect of leukemia-initiation (**Fig. 2e-g**) and a ∼40-fold reduction of leukemia-initiating cells (LIC; 1 in 1,031 to 1 in 40,055 cells), which suggests that Lgr5 is crucial for B-ALL leukemia-initiation. We next stained patient-derived B-ALL xenograft cells (PDX2) for LGR5 surface expression, sorted LGR5^+^ and LGR5^-^ cells by flow cytometry and compared leukemia-initiation from 100,000, 10,000 and 1,000 injected cells (**Extended data figure 7f, g**). Compared to LGR5^+^ B-ALL cells, development of leukemia was delayed when LGR5^-^ B-ALL cells were injected at low cell numbers, however both populations were able to initiate fatal leukemia. Interestingly, regardless of whether LGR5^+^ or LGR5^-^ B-ALL cells were initially injected, B-ALL cells gave rise to mixed populations that contained similar fractions of LGR5^+^ and LGR5^-^ B-ALL cells, (**Extended data figure 7**). Sorting and clonal outgrowth from single cells confirmed that LGR5^-^ B-ALL cells can give rise to LGR5^+^ progeny and *vice versa* (**Extended data figure 7d**). These results suggest that LGR5 is required for LIC function in B-ALL but not a surface marker that would determine a permanent stem cell hierarchy. The single-cell sort experiment showed that most, if not all B-ALL cells have the capacity to express LGR5 and transiently acquire LIC-potential (**Extended data figure 7d**). Forced expression of LGR5 increased clonal competitiveness (**Fig. 2h**) and the ability to form colonies in patient-derived B-ALL xenografts (**Extended data figure 7a**) and murine B-ALL cells (**Fig. 2i**). A limiting-dilution transplantation experiment with patient-derived PDX2 B-ALL cells revealed that forced expression of LGR5 increased LIC-potential **(Fig. 2j, Extended data figure 7b-c**) and the frequency of LIC by ∼8-fold (from 1 in 821 to 1 in 97; **Fig. 2j**).

**Figure 6:**
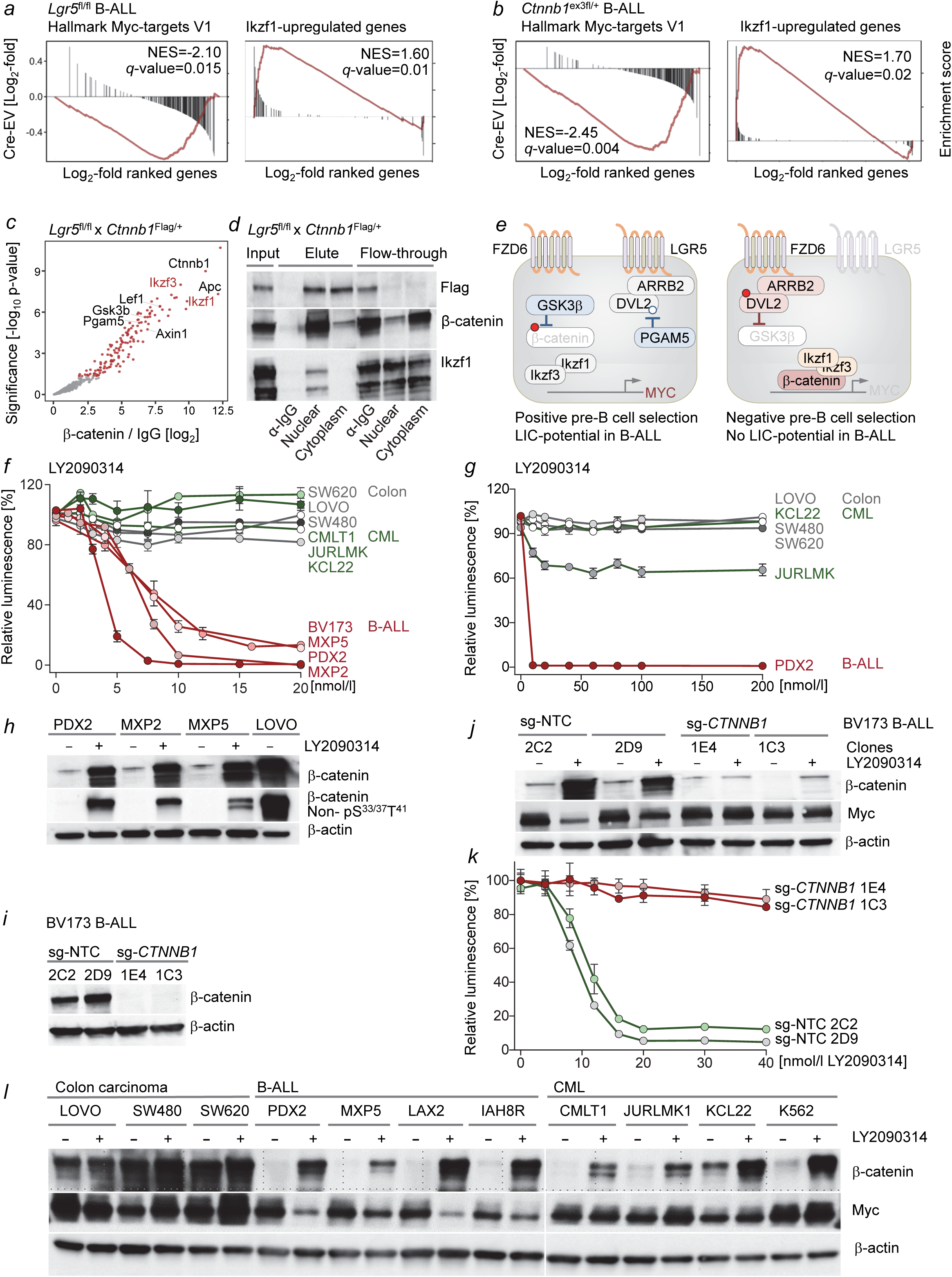
Pharmacological hyperactivation of β-catenin signaling represents a strategy to eradicate B-ALL. BCR-ABL1-driven *Lgr5*^fl/fl^ (**a**) and *Ctnnb1*^ex3fl/+^ (**b**) B-ALL cells were transduced with 4-OHT-inducible Cre-ER^T2^ or ER^T2^. Gene expression changes following 4-OHT-treatment were studied by RNA-seq analysis (**Fig. 3b**). Gene set enrichment analysis showed depletion of MYC target gene expression defined by Hallmark-MYC Targets V1 but increases of IKZF1-upregulated gene sets. To elucidate why β-catenin-activation results in transcriptional repression of Myc in pre-B cells, unlike other cell types, we transformed pre-B cells from Lgr5^fl/fl^ x *Ctnnb1*^flag/+^ knockin mice that express FLAG-tagged β-catenin. Upon Cre-mediated deletion of *Lgr5*, we identified binding partners by pull-down of FLAG-tagged β-catenin and proteomic analyses (**c**). Pull-down with anti-IgG antibody was performed to normalize for non-specific binding. Proteins bound to β-catenin and IgG antibody were analyzed by mass spectrometry and are plotted based on significance and log2-fold enrichment over IgG (n=4). Proteins with the highest nuclear enrichment of binding to β-catenin included Ikzf1 (Ikaros) and Ikzf3 (Aiolos) that function as transcriptional repressors of Myc in pre-B cells (highlighted in red). The β-catenin-Ikzf1 interaction was confirmed by co-immunoprecipitation and Western blot (**d**), using antibodies to detect FLAG, β-catenin and Ikzf1 in whole cell lysates (input), proteins bound (elute) and not bound (flow-through) by IgG or β-catenin antibodies (**d**). Unlike colon cancer and CML, β-catenin associates with Ikzf1 and Ikzf3 in pre-B cells, which could explain that β-catenin accumulation (e.g. as the result of loss of *Lgr5*) results in transcriptional repression of Myc in pre-B cells as opposed to transcriptional activation in other cell types (**e**). B-ALL cells (red), CML (green) and colon cancer (gray) cell lines were treated with the GSK3β inhibitor LY2090314 at concentrations of (**f**) 0 up to 20 nM or (**g**) 0 up to 200 nM for 3 days. and cell viability was determined. All B-ALL cells expressed functional IKZF3. BV173 and MXP5 carry *IKZF1* deletions. (**h**) B-ALL PDX and colon cancer (LOVO) cells were treated with the GSK3β inhibitor LY2090314 for 16 hours. Western blot analyses were performed for β-catenin, stable β-catenin lacking GSK3β-phosphorylation (S^33^, S^37^, T^41^) and β-actin. (**i**) BV173 B-ALL cells were electroporated with Cas9-RNPs with non-targeting crRNAs (sg-NTC) or crRNAs targeting β-catenin (sg-*CTNNB1*). Single cell-derived colonies were generated, expanded in cell culture. Deletion of β-catenin was confirmed in two clones (1E4, 1C3) by Western blot (**i**). BV173 B-ALL cells (*IKZF1*-deletion) with (sg-*CTNNB1*) or without (sg-NTC) deletion of β-catenin were treated for 12 hours with LY2090314 (20 nM) and Western blot analyses for β-catenin and Myc levels were performed using β-actin as loading control (**j**). B-ALL cells with (clones 1E4, 1C3; sg-*CTNNB1*) or without (clones 2C2, 2D9; sg-NTC) deletion of β-catenin were treated with LY2090314 for 3 days at the indicated concentrations and relative viability was determined by luminescence measurements (**k**). Colon cancer, B-ALL and CML cells were treated with LY2090314 at 10 nM for 12 hours and β-catenin and Myc levels were measured in these cells by Western blot, using β-actin as loading control (**l**).

## Stemness in pre-B cells and B-ALL is tied to positive selection and transient expression of LGR5

While stemness and LIC-capabilities in myeloid leukemia and solid tumors follow a developmental hierarchy^27-29^, these results suggest a transient model of stemness in B-ALL that is tied to positive selection and short-lived expression of LGR5, comparable to the narrow window of LGR5 expression at the pre-BCR checkpoint during normal B-cell development. Self-renewal in pre-B-cells is determined by positive selection based on their ability to express a functional pre-BCR, which then signals transient activation of Lgr5. A static, developmentally-determined hierarchy with a rare primitive stem cell at its apex, as in hematopoietic stem cells^30^, would broadly enable developmental progression but not positive selection of the few precursor cells that have productively rearranged Ig V, D and J segments to express a functional pre-BCR. Unlike myeloid and epithelial cells, we propose that self-renewal in early B-cell development is not a pre-determined fate but driven by positive selection signals from a functional pre-BCR. In B-ALL, self-renewal and positive selection are induced by oncogenic mimics of pre-BCR-signaling^31-36^. Hence, stemness and LIC-potential in B-ALL are tied to oncogenic signals that mimic positive selection signals from a functional pre-BCR. This scenario would reconcile the perplexing finding that unlike rare and phenotypically primitive stem cell populations in myeloid malignancies and solid tumors, LIC in B-ALL are very frequent and do not conform to a developmentally determined stem cell hierarchy^37-40^.

## Lgr5 functions as a negative regulator of β-catenin

To elucidate the mechanism of Lgr5-dependent leukemia-initiation and LIC survival in B-ALL, we assessed the consequences of *Lgr5*-deletion at the level of signal transduction. Lgr5 belongs to the rhodopsin-like class-A (subfamily 10) of G-protein coupled receptors, which signal through the G-protein-coupled receptor kinase 2 (GRK2) in B-cells^41^. Hence, we first performed a global proteomic analysis of phosphorylation changes upon inducible *Lgr5*-deletion in B-ALL cells. Consistent with a role of LGR5 in the regulation of WNT/β-catenin-signaling, the most prominent phosphorylation-changes changes affected Axin 2, a negative regulator of β-catenin^42^ and β-catenin itself (S^675^; **Fig. 3a**). The phosphorylation on β-catenin-S^675^ is mediated by cAMP-dependent protein kinase (PKA) and results in increased β-catenin activity^43^, which was unexpected given the known role of LGR5 in positive regulation of WNT signaling in epithelial stem cells^12^. Other PKA-related molecules that were phosphorylated upon *Lgr5*-deletion included Akap12 and Pde8a, whereas phosphorylation of Myc and its target E2f2 was decreased (**Fig. 3a**). Measuring the impact of *Lgr5*-deletion at the transcriptional level, RNA-seq analysis revealed upregulation of multiple surface receptors that are associated with anergy and exhaustion of B- and T-cells, including Ctla4, Tigit, Prdm1, Ccr2, Dgka, Cd5, Il2ra (Cd25) and Cd244 (2B4). These changes were consistent with a recently identified signature of B-cell anergy and negative selection that depends on the Ikzf1 transcription factor^44^. The PKA regulatory subunit Prkar1b and PKA-scaffold Akap12 **(Fig. 1i**; **Extended data figure 1d**) were upregulated upon *Lgr5*-deletion, whereas Myc and E2f2 were downregulated (**Fig. 3b**). Western blot analyses of *Lgr5*-deletion in B-ALL cells confirmed upregulation of Dgka, Akap12, Cdkn2a, Prdm1 and Zap70 (**Fig. 3c**; **Extended data figure 8a**), while upregulation of anergy- and exhaustion markers (Cd5, Il2ra, Ccr2, Ctla4, Cd244) was confirmed by flow cytometry (**Fig. 3i**). Strikingly, *Lgr5*-deletion resulted in substantial depletion of Axin2 protein, which was paralleled by >250-fold accumulation of β-catenin (**Fig. 3c**). Upregulation of Axin2 mRNA upon Lgr5-deletion is seemingly in contrast with loss of Axin2 protein. However, *Axin2* represents a classical WNT/β-catenin target gene^45^, hence massive accumulation of β-catenin would be expected to increase transcriptional activation of Axin2. Consistent with loss of Axin2 and increased β-catenin stability, phosphorylation of β-catenin serine (S^33^, S^37^) and threonine (T^41^) residues was reduced (**Fig. 3c**). The sites in β-catenin exon 3 are phosphorylated by GSK3β and initiate degradation. In addition, PKA-mediated β-catenin phosphorylation of S^675^ identified by phosphoproteomic analyses (**Fig. 3a**) was confirmed by Western blot (**Fig. 3 c**). These results suggest a central role of LGR5 in negative regulation of WNT/β-catenin signaling in pre-B cells and B-ALL.

## Lgr5-dependent sequestration of Dishevelled2 enables GSK3β-mediated negative regulation of β-catenin

To identify LGR5-interaction partners that could explain the underlying mechanism of LGR5 function and negative regulation of β-catenin in B-ALL cells, we generated C-terminal fusion proteins between LGR5 and the bacterial biotin-ligase BirA (**Extended data figure 8**). Proximity-based analyses of proteins that were biotinylated by LGR5-BirA fusions (Bio-ID) identified Dishevelled2 (DVL2), its scaffold β-arrestin (ARRB2)^14, 17, 19^ and the DVL2 clathrin adapter AP2 (AAK1, AP2A2, AP2B1)^15^ as central interaction partners of the cytoplasmic tail of LGR5 (**Fig. 3d**). The ability of DVL2 to bind to the cytoplasmic tail of FZD receptors is critical for WNT/β-catenin signaling^46^. When bound to FZDs, DVL2 is heavily phosphorylated and functions as a negative regulator of GSK3β (**Extended data figure 8e-f**)^42^, hence promoting β-catenin stabilization. Interestingly, the LGR5 Bio-ID analysis also identified the phosphatase PGAM5, which negatively regulates β-catenin by dephosphorylation of DVL2^16, 48^. Binding of DLV2, its scaffolds ARRB2, AP2B1 and AAK1 as well as the DLV2-phosphatase PGAM5 to the LGR5 tail was confirmed by pull-down experiments using LGR5 with a C-terminal HA-tag (**Extended data figure 8d**). Based on these observations, we propose that LGR5 exerts negative regulation of β-catenin via sequestration of DVL2 and its scaffolds ARRB2 and AP2 from FZD receptors (**Extended data figure 8e-f**). According to this scenario, FZD and LGR5 are structurally similar GPCRs that compete for binding to DVL2. In positively selected pre-B cells and B-ALL cells, high expression levels of LGR5 lead to sequestration of DVL2 from FZD receptors. In proximity to LGR5, DVL2 is dephosphorylated by PGAM5^48^, which relieves DVL2-mediated inhibition of GSK3β^47^, thereby allowing GSK3β to phosphorylate β-catenin for subsequent degradation. Upon *Lgr5*-deletion, DVL2 associates with FZD receptors. Bound to FZDs, DVL2 is constitutively phosphorylated^46^, inhibits GSK3β, which results in accumulation of β-catenin and activation of WNT-signaling^46^ (**Extended data figure 8f**).

## β-catenin-hyperactivation represents the functional equivalent of Lgr5-deletion

If a main function of LGR5 was to limit β-catenin activity, one would predict that inducible ablation of *Lgr5* recapitulates the transcriptional program of hyperactive WNT/β-catenin signaling. To test this hypothesis, we generated BCR-ABL1-driven B-ALL cells from mouse *Ctnnb1*^ex3fl/+^ bone marrow pre-B cells. Cre-mediated excision of exon 3 of β-catenin in this mouse model removes GSK3β phosphorylation sites (S^33^, S^37^ and T^41^) for β-catenin degradation (Harada et.al. 1999). Hence, activation of Cre in *Ctnnb1*^ex3fl/+^ B-ALL cells caused stabilization and accumulation of β-catenin (**Extended data figure 9**). Gene expression changes resulting from β-catenin hyperactivation mirrored multiple features of *Lgr5*-deletion including downregulation of Myc (**Fig. 3e-f**) and E2f2 (**Fig. 3f**) and induction of B-cell anergy phenotypes (Ctla4, Prdm1, Ccr2, Dgka, Cd5, Il2ra and Cd244; flow cytometry validation **Fig. 3i-j**). Gene set enrichment analysis revealed a striking global similarity of *Ctnnb1*^ex3fl/+^ and *Lgr5*^fl/fl^ transcriptomes in B-ALL cells **(Fig. 3g-h**), suggesting that β-catenin-hyperactivtion represents the functional equivalent of *Lgr5*-deletion at the transcriptional level.

## Lgr5-mediated negative regulation of β-catenin is essential for B-lymphopoiesis and B-ALL initiation

Like inducible deletion of *Lgr5* in pre-B and B-ALL cells, also inducible hyperactivation of β-catenin (*Ctnnb1*^ex3fl/+^) subverted competitive fitness (**Fig. 4a, Extended data figure 9a-c**), reduced colony forming capacity (**Fig. 4b**) and increased the fraction of cells in G0/G1 phase of the cell cycle (**Fig. 4c**). Interestingly, however, despite these defects, neither *Lgr5* -deletion nor hyperactivation of β-catenin substantially increased apoptosis or cell death (**Extended data figure 9d-g**). These findings suggest that Lgr5 -and its ability to curb β-catenin activation-are essential for self-renewal and clonal fitness of B-ALL cells, but not for survival. To experimentally test whether negative regulation of β-catenin is the mechanistic basis of Lgr5 function during early B-lymphopoiesis and B-ALL initiation, we tested whether genetic deletion of β-catenin (*Ctnnb1*; **Extended data figure 10**) can rescue the deleterious effects of *Lgr5*-ablation in B-ALL cells. While β-catenin serves critical functions in embryonic development and multiple cell lineages^49,50^. *Ctnnb1*-deletion did not have major impact on normal hematopoiesis^51^. Likewise, *Ctnnb1*-deletion did not affect *Lgr5*^fl/fl^ B-ALL cells in our analysis. However, when Cre was induced for concurrent *Lgr5*-deletion, loss of *Ctnnb1* largely rescued clonal fitness in a competitive cell culture assay (**Fig. 4d**). Strikingly, deletion of *Ctnnb1* entirely restored Myc-expression (**Fig. 3e**), colony formation ability (**Fig. 4e**) and proliferation (**Fig. 4f**). Together with the finding that *Lgr5*-deletion recapitulates the transcriptional program of β-catenin hyperactivation (**Fig. 3f-h**), these experiments provide genetic evidence that Lgr5-mediated restraint of β-catenin is central to its function during B-lymphopoiesis and B-ALL initiation.

## Negative regulation of β-catenin by Lgr5 is limited to early B-cell development

Studying the entire spectrum of hematopoietic populations, Lgr5 expression was found exclusively at the pre-BCR checkpoint during early B-cell development (**Fig. 1a-c**; **Extended data figure 1b-c**). This extremely narrow window of gene expression was mirrored by the specific requirement of Lgr5 at the pre-BCR checkpoint: deletion of *Lgr5* in mature and GC-B-cells had only very subtle effects **(Extended data figure 3-4**) whereas deletion prior to the pre-BCR checkpoint resulted in complete loss of B-cell production (**Fig. 1f, Extended data figure 2**). Strikingly, deletion of *Lgr5* in pre-B cells caused a >250-fold increase of β-catenin protein levels (**Fig. 3c, Fig. 4g**) but had no measurable effect on β-catenin levels in mature B-cells (**Fig. 4g**). Consistent with the identification of negative regulation of β-catenin as central mechanism of Lgr5 function, these results corroborate that mature B-cells no longer depend on Lgr5 function (**Fig. 4h**).

## Negative regulation of β-catenin in B-cell malignancies

Oncogenic WNT/β-catenin signaling and activating mutations that increase stability and transcriptional activity of β-catenin represent a common feature throughout all main types of cancer^7, 8^. In a comprehensive analysis of 66,820 samples encompassing 15 types of cancer (COSMIC)^52^, we identified 4,971 activating *CTNNB1* mutations (on average 7.4%). Strikingly, however, in 2,375 B-cell malignancies, no activating *CTNNB1* mutations were found (expected 176, observed 0, χ^²^ test *P*=0.0001; **Fig. 5a**). Likewise, an analysis of β-catenin expression at the protein level (Protein Atlas)^53^ revealed frequent activation of WNT/β-catenin signaling in cancer but not in B-cell malignancies. 519 tumor samples representing 17 types of cancer were stained for β-catenin expression using three different antibodies (1,521 experiments total) and analyzed by immunohistochemistry. Among 1,488 solid tumor samples, in 1,202 cases (81%) β-catenin expression was detected, as compared to none in 33 B-cell lymphoma samples (expected 26.7, observed 0, χ^²^ test *P*=0.0001; **Fig. 5b**). These results were further corroborated by the analysis of β-catenin expression in a reverse phase protein array data set (CCLE)^54^ encompassing 957 cancer cell lines. tSNE plots based on RPPA data for 214 proteins segregated the 110 B-cell tumor cell lines from the 847 cancer cell lines (**Fig. 5c**) and revealed strikingly low β-catenin RPPA signals for the 110 B-cell tumor cell lines, including 20 B-ALL, compared to cancer cell lines (**Fig. 5d**). While these results were based on publicly accessible data sets^52-54^, we confirmed lack of oncogenic β-catenin signaling and expression in B-cell malignancies by Western blot and immunohistochemistry (**Fig. 5e-f**; **Extended data figure 11a**). By nuclear and cytoplasmic fractionation and Western blot analysis, we consistently detected nuclear β-catenin in solid tumors, including non-small cell lung cancer (NSCLC), colon cancer and malignant melanoma (n=16), but not in B-ALL (n=7) and mature B-cell malignancies (n=15; **Fig. 5e**; **Extended data figure 11a**). Likewise, strong expression of β-catenin was observed by immunohistochemistry in normal colon tissue, colon cancer, melanoma, NSCLC and breast cancer, but not in normal lymphoid follicles (tonsil) and B-cell lymphoma (**Fig. 5f**). Based on *CTNNB1* mutation data, immunohistochemistry of β-catenin expression, RPPA data and our own corroborating experiments, these results highlight that B-cell malignancies differ from other types of cancer in that they are consistently exempt from oncogenic activation of the WNT/β-catenin pathway.

## Lgr5 function in B-ALL compared to colon cancer cells

While not interchangeable, Wnt ligands (e.g. Wnt3a) of FZD receptors and R-spondin ligands (e.g. Rspo1) of Lgr5 cooperate to potentiate Wnt/β-catenin signaling. Rspo1 is thought to act as ligand for Lgr5^12, 18^, which then counteracts the RNF43 and ZNRF3 E3 ubiquitin ligases to stabilize Fzd receptors for prolonged Wnt/β-catenin signaling. Based on recent evidence, however, Rspo2 can directly inhibit RNF43 and ZNRF3 and promote Wnt/β-catenin signaling even in the absence of Lgr5^56^. In B-ALL cells, deletion of *Lgr5* resulted in a >250-fold accumulation of nuclear β-catenin, however, Wnt3a and Rspo1 had no measurable effect on β-catenin levels, neither alone nor in combination (**Extended data figure 11b**). Since deletion of *Lgr5* caused massive accumulation of nuclear β-catenin and loss of self-renewal capacity in B-ALL cells, we tested whether *LGR5*-deletion in human colon cancer cells (LOVO) had similar effects. Deletion of *LGR5* in LOVO cells was achieved by electroporation-based delivery of Cas9 ribonucleoproteins (RNPs) containing Cas9 and guide-RNAs directed against *LGR5* or a non-targeting control and subsequent screen of single clones for loss of *LGR5* (**Extended data figure 11c**). Unlike B-ALL, however, genetic loss of *LGR5* neither affected outgrowth from single clones nor β-catenin levels. Hence, LGR5-mediated negative regulation of β-catenin is likely a unique feature of pre-B-cells. Conversely, Wnt and Rspo ligands are important regulators of β-catenin in epithelial cell types but have no apparent function in pre-B and B-ALL cells.

## Contribution of PKA and RHOA signaling to LGR5-function in B-ALL cells

Given the nature of Lgr5 as a G-protein coupled receptor and its association with Grk2, PKA subunits (Prkacb), PKA-adapters (Akap12, Akap8, Akap1; **Fig. 3d**; **Extended data figure 8d**) and LGR5-dependent downstream phosphorylation changes of Rock2, Gnai2 (Gαi2) and PKA-associated Akap12 and Pde8a (**Fig. 3a**) as well as PKA-mediated phosphorylation of β-catenin (S^675^; **Fig. 3a**), we examined potential contributions of the PKA-pathway to LGR5 function. Inducible Cre-mediated deletion of *Lgr5* in B-ALL induced significant increases of PKA activity (**Extended data figure 12a**). To determine whether hyperactivation of PKA signaling contributes to the deleterious effects of *Lgr5*-ablation, we performed rescue experiments based on deletion of PKA subunits. These deletions were introduced into *Lgr5*^fl/fl^ B-ALL cells by retroviral delivery of Cas9 and guide-RNAs directed against *Prkaca* and *Prkacb* or a non-targeting control (NTC; **Extended data figure 12**). While genetic deletion of *Ctnnb1* almost entirely rescued the deleterious effects of *Lgr5*-deletion, deletion of *Prkaca* and *Prkacb* partially restored competitive fitness and colony formation (**Extended data figure 12c-f**), consistent with increased phosphorylation of the activating PKA-site S^675^ on β-catenin (**Fig. 3a, c**)^43^. Like PKA, enzymatic activity of RHOA, upstream of Rock2 and Gαi2 (**Fig. 3a**) was strongly upregulated upon Cre-mediated deletion of *Lgr5* in B-ALL cells (**Extended data figure 13a**). Since Rhoa functions as a downstream target of FZD-DVL2 signaling^57^, we tested whether Cas9-mediated deletion of *Rhoa* was able to restore competitive fitness and colony formation in B-ALL cells. However, unlike deletion of *Ctnnb1* and *Prkaca*-*Prkacb*, loss of *Rhoa* had no mitigating effect on *Lgr5*-deletion (**Extended data figure 13b-e**).

## β-catenin forms complexes with Ikzf1 and Ikzf3 in pre-B cells for transcriptional repression of Myc

Decreased phosphorylation of E2f2 (S^314^) and Myc (T^79^; **Fig. 3a**) and transcriptional downregulation of E2f2 and Myc was identified as a common feature of *Lgr5*-deletion (**Fig. 3b**) and direct β-catenin hyperactivation (**Fig. 3f**). Consistent with a major role of Myc downstream of Lgr5 and β-catenin, we found that Myc transcriptional targets were globally suppressed both upon deletion of *Lgr5* (*Lgr5*^fl/fl^; **Fig. 6a**) and β-catenin hyperactivation (*Ctnnb1*^ex3fl/+^; **Fig. 6b**). For this reason, we tested whether reconstitution of Myc could rescue the deleterious effects of *Lgr5* deletion and β-catenin hyperactivation. Myc overexpression restored competitive fitness of B-ALL cells that was lost upon *Lgr5*-deletion (**Extended data figure 14a, c**) or β-catenin hyperactivation (**Extended data figure 14b, d**). Indeedx, both *Lgr5*-deletion and inducible stabilization of β-catenin caused profound suppression of Myc in B-ALL cells (**Fig. 3e**; **Extended data figure 14**), which is in striking contrast to Myc as a classical target of β-catenin-mediated transcriptional activation^9^. To elucidate why β-catenin functions as transcriptional repressor of Myc in pre-B cells, unlike other cell types, we identified β-catenin interacting proteins in B-ALL cells. To this end, we transformed pre-B cells from *Mb1*^Cre-ERT2^ x *Lgr5*^fl/fl^ x *Ctnnb1*^flag/+^ knockin mice that express FLAG-tagged β-catenin from one allele. Upon Cre-mediated deletion of Lgr5, we identified binding partners by pull-down of FLAG-tagged β-catenin and proteomic analyses (**Fig 6c**; **Extended data figure 15a**). Surprisingly, the proteins with the highest nuclear enrichment of binding to β-catenin included Ikzf1 (Ikaros) and Ikzf3 (Aiolos), two pre-B cell-specific transcription factors that function as transcriptional repressors of Myc^10^. The interaction between β-catenin and Ikzf1 was confirmed by co-immunoprecipitation and Western blot (**Fig. 6c**). We propose that that β-catenin binds to the *MYC* promoter in pre-B cells, as in other cell types. Unlike β-catenin/TCF4 complexes in other cell types^7, 9^, however, β-catenin associates in pre-B cells with Ikzf1 and Ikzf3. Given that both Ikzf1 and Ikzf3 are specifically active at the pre-BCR checkpoint^10, 11^, the discovery of nuclear complexes between β-catenin and Ikzf1-Ikzf3 in pre-B cells could explain that β-catenin functions as transcriptional repressor of Myc in pre-B and B-ALL (**Fig. 6e**) as opposed to transcriptional activation in other cell types. Besides negative regulation of Myc, β-catenin-Ikzf1 complexes more broadly promoted an Ikzf1-dependent transcriptional program (**Extended data figure 15b-c**), that recapitulates multiple features of Ikzf1-dependent transcriptional programs of B-cell anergy and negative selection^44^.

## Therapeutic targeting of Lgr5^+^ leukemia-initiating cells (LIC) in B-ALL

Given that Lgr5 serves a critical role in self-renewal and survival of LIC in B-ALL **(Fig. 2a-j**), we tested whether eradication of Lgr5^+^ cells may be useful for therapeutic purposes. To eradicate cancer stem cells in colon cancer, an antibody-drug conjugate (ADC) targeting LGR5 was recently developed and demonstrated potent efficacy and safety *in vivo*^24^. Since Lgr5 mRNA levels in B-ALL cells are similar or even higher compared to colon cancer (**Fig. 1g**), we tested the efficacy of LGR5-ADC in a xenotransplantation model based on B-ALL PDX. NSG mice bearing B-ALL PDX were treated with anti-LGR5 (8E11) conjugated to a potent microtubule inhibitor, monomethyl auristatin E (MMAE) or an isotype control antibody (anti-gD) linked to MMAE. Compared to anti-gD-vc-MMAE, anti-LGR5-vc-MMAE substantially reduced leukemia burden and significantly extended overall survival (*P*=0.028; **Extended data figure 16a-b**). However, when treatment was stopped (day 50), mice recipients ultimately relapsed and died from B-ALL. These findings suggest that repeated injections with anti-LGR5-vc-MMAE achieved long-term disease control but not complete LIC-eradication.

## Pharmacological hyperactivation of β-catenin leverages a unique vulnerability in B-ALL cells

Our mechanistic experiments revealed that hyperactivation of β-catenin and β-catenin-mediated transcriptional repression of MYC represent the underlying basis of the unexpected dependency of B-ALL cells on LGR5. Hence, as an alternative to LGR5-ADC mediated targeting of LGR5^+^ B-ALL cells, we tested pharmacological hyperactivation of β-catenin, and, presumably, β-catenin-mediated repression of *MYC*, to eradicate LIC in patient-derived B-ALL. To this end, we tested two FDA-approved small molecule inhibitors of GSK3β (9-ING-41 and LY2090314; **Extended data figure 16**) that have demonstrated safety in clinical trials for patients with metastatic cancer and acute myeloid leukemia (AML)^58^. Inhibition of GSK3β would be expected to increase stability of β-catenin in a similar way as deletion of the GSK3β-phosphorylation sites (exon 3) in the *Ctnnb1*^ex3fl/+^ mouse model (**Fig. 3e, Extended data figure 9a**)^59^. Strikingly, LY2090314 had profound effects on B-ALL cells at low nanomolar concentrations but not colon cancer or myeloid leukemia cells (**Fig. 6f-g**). Even at >40-fold higher concentrations, LY2090314 had no substantial toxic effects on colon cancer and myeloid leukemia cells (**Fig. 6g**). The toxic effect on B-ALL cells was paralleled by massive accumulation of β-catenin, comparable to baseline levels in colon cancer cells (**Fig. 6h, l**). While structurally similar to LY2090314 (**Extended data figure 16e**), 9-ING-41 was not selective for B-ALL and killed B-ALL, colon cancer and myeloid leukemia cells only at much higher concentrations. Unlike LY2090314, 9-ING-41 failed to increase β-catenin protein levels in B-ALL. In colon cancer cells, β-catenin levels were >250-fold higher than in B-ALL cells, but remained unchanged after treatment with 9-ING-41. To test our hypothesis that accumulation of β-catenin represents the mechanistic basis of LY2090314-mediated toxicity in B-ALL cells, we deleted *CTNNB1* in the human B-ALL cell line BV173 using Cas9-RNPs and screening of clones for *CTNNB1*-deletion from single cells. Interestingly, deletion of *CTNNB1* conferred complete resistance of B-ALL cells to LY2090314. Importantly, preventing β-catenin upregulation in response to LY2090314-treatment also mitigated suppression of Myc, which would otherwise result from LY2090314-induced β-catenin hyperactivation (**Fig. 6i-k**). Further corroborating β-catenin-induced repression of Myc as the underlying mechanism of LY2090314-induced toxicity in B-ALL cells, we tested the effect of LY2090314 on β-catenin and Myc levels in resistant colon cancer and myeloid leukemia cells as well as sensitive B-ALL cells. β-catenin levels were constitutively high in colon cancer and did not increase upon drug-treatment. Resistant myeloid leukemia cells responded to LY2090314 by strong upregulation of β-catenin (**Fig. 6l**). However, unlike B-ALL cells, massive accumulation of β-catenin did not come at the expense of Myc, presumably because myeloid cells lack expression of IKZF1 and IKZF3.

## IKZF1 and IKZF3 set the threshold for β-catenin-induced transcriptional repression of MYC

These pre-B cell-specific transcription factors function as repressors of MYC^10^ and associate with β-catenin in B-ALL cells (**Fig. 6e, 6l**; **Extended data figure 15**). Likewise, colon cancer cells, lacking expression of IKZF1 and IKZF3, expressed β-catenin at >250-fold higher baseline levels than B-ALL cells, but without any apparent impact on expression levels of MYC. β-catenin/TCF4 complexes drive transcriptional activation of MYC^7, 9^ in many cell types, including colon cancer and CML cells. However, β-catenin associated with the B-lymphoid transcription factors Ikzf1 and Ikzf3 has the opposite function in pre-B cells and functions as a powerful transcriptional repressor of Myc. Hence, Ikzf1 and Ikzf3 compete with TCF4 for binding to β-catenin and thereby determine the outcome of β-catenin signaling, e.g. transcriptional activation (TCF4) or repression (Ikzf1, Ikzf3) of MYC. Importantly, transcriptional repression of MYC by IKZF1 and IKZF3 is restricted to pre-B cells^10^, which explains the narrow window of LGR5 expression and sensitivity to β-catenin hyperactivation at the pre-BCR checkpoint. For instance, deletion of *Lgr5* and β-catenin hyperactivation did not affect mature B-cells (**Fig. 4g-h**; **Extended data figure 3,4**). IKZF1 and IKZF3 are essential for oncogenic signaling and therapeutic targets in multiple myeloma^60^, a B-lymphoid malignancy derived from terminally differentiated B-cells. In contrast, both *IKZF1* and *IKZF3*^61^ are tumor suppressors and frequently deleted in human B-ALL. Importantly, multiple oncogenic drivers in B-ALL, including TCF3-PBX1^62^ and BCR-ABL1^63^ can activate β-catenin for increased proliferation and survival. Hence, deletion of *IKZF1* or *IKZF3* may allow for increased activation of β-catenin in B-ALL cells, while LGR5 functions as feedback regulator to buffer fluctuations of β-catenin signaling. Importantly, *IKZF1*-deletion represents a particularly impactful predictor of poor clinical outcomes in patients with B-ALL^64^. However, patients with *IKZF1*-deletion retain *IKZF3* function and deletion of the two transcription factors is mutually exclusive^61^. In our analysis, both MXP5 and BV173 B-ALL cells harbored *IKZF1* deletions. However, despite IKZF1-deletion, B-ALL remained fully sensitive to GSK3β-inhibition (**Fig. 6f**; **Extended data figure 14e**) and downregulated MYC expression upon treatment with LY2090314 (**Fig. 6j, 6l**). While these responses were slightly diminished compared to B-ALL samples without *IKZF1*-deletion, these results suggest that IKZF3 alone is sufficient to repress MYC and induce cell death. Hence, acute hyperactivation of β-catenin (e.g. by GSK3β-inhibition, LY2090314) likely represents a powerful therapeutic approach even for high-risk B-ALL cases with *IKZF1*-deletion.

## Concluding remarks

Previous studies demonstrated that hyperactivation of β-catenin can have deleterious consequences in the hematopoietic system^65, 66^ and that multiple mechanisms, e.g. LMBR1L-mediated ubiquitination of FZD6 and LRP6^67^ are necessary to achieve optimal dosage and prevent toxic levels of β-catenin hyperactivation. Here we show that LGR5-mediated sequestration of DVL2 is essential to curb β-catenin activation at the pre-BCR checkpoint. Pre-B cells that fail to express a functional pre-BCR do not upregulate Lgr5. In the absence of Lgr5, uncontrolled β-catenin accumulation engages an Ikzf1-dependent transcriptional program of negative pre-B cell selection and anergy^44^ resulting in clonal deletion. Positively selected Lgr5^+^ pre-B cells retain expression of MYC and initiate clonal expansion. Thereby, Lgr5 prevents β-catenin accumulation and formation of β-catenin complexes with IKZF1 and IKZF3 for transcriptional repression of MYC. While this mechanism is critical for the initiation of normal B-lymphopoiesis, it also enables leukemia-initiation in B-ALL. As shown by treatment studies with LGR5-ADC and GSK3β small molecule inhibitors, this unexpected vulnerability can be leveraged for targeted LIC-eradication in drug-resistant B-ALL.

## Acknowledgments

We would like to thank Dr. Michael Kahn, Department of Molecular Medicine, COHCCC, for critical discussions, Lars Klemm and Dr. Anna Hecht for their help with technical aspects of experiments and current and former members of the Müschen laboratory for their support and helpful discussions. Research in the Müschen laboratory is funded by the NIH through an NCI Outstanding Investigator Award R35CA197628 (to M.M.), U10CA180827 (to M.M.), R01CA137060, R01CA157644, R01CA172558 and R01CA213138 (to M.M.), the Howard Hughes Medical Institute HHMI-55108547 (to M.M.), the Norman and Sadie Lee Foundation (to M.M.), and the Falk Trust through a Falk Medical Research Trust Catalyst Award (to M.M.), the Pediatric Cancer Research Foundation (PCRF), and the California Institute for Regenerative Medicine (CIRM) through DISC2-10061. M.M. is a Howard Hughes Medical Institute (HHMI) Faculty Scholar.

