## Extended Data Figures for "Lgr5-mediated restraint of β-catenin is essential for B-lymphopoiesis and leukemia-initiation"

**Extended data figure 1:** *Identification of Lgr5 and Akap12 expression at the pre-BCR checkpoint*

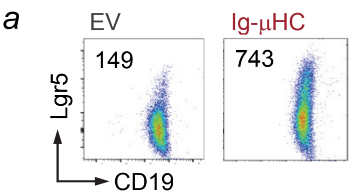


(**a**) *Rag2*$^{-/-}$ pro-B cells were transduced with a functional 4-OHT-inducible Ig µ-heavy chain (µHC) to initiate pre-BCR signaling. Representative FACS analysis of Lgr5 surface levels 24 hours after pre-BCR induction with mean fluorescence intensity (MFI) values is shown.

(**b**) *Lgr5* mRNA levels (red color scale) as determined by RNA-seq were plotted for hematopoietic lineages (B-lymphoid, T-lymphoid, NK-cell, myeloid and erythroid) as well as differentiation stages within each lineage as dendrogram (**b**) and bar chart (**c**).

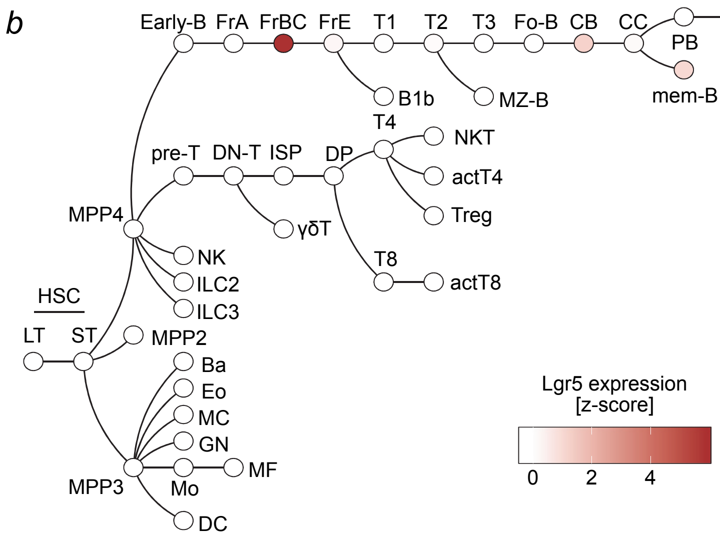


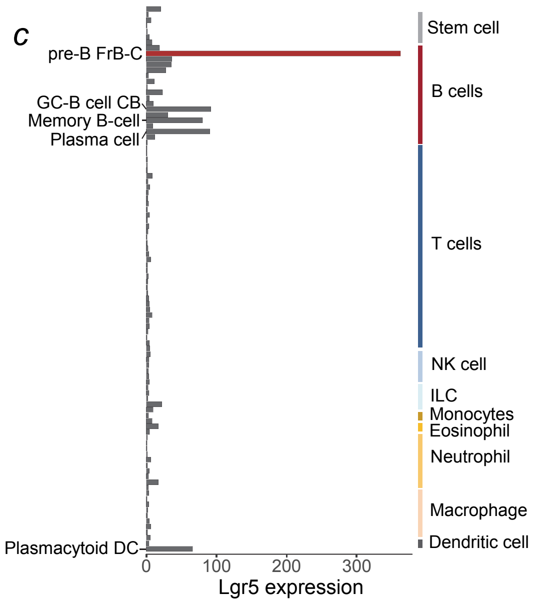


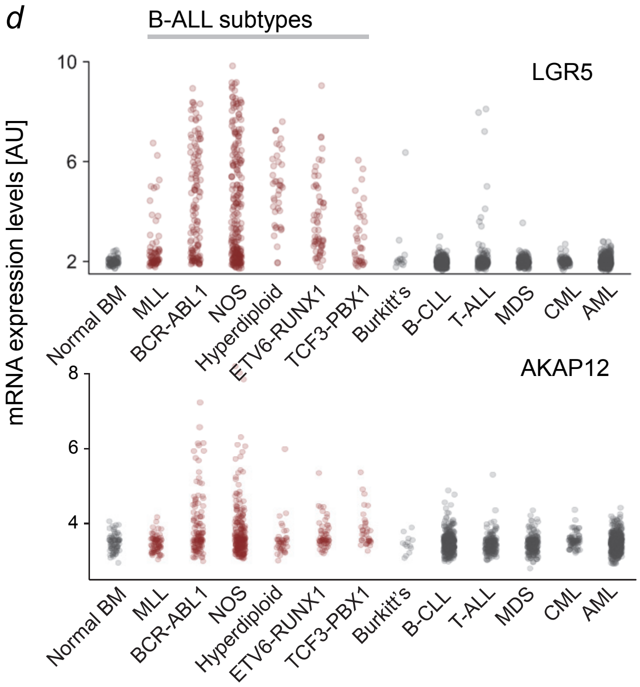


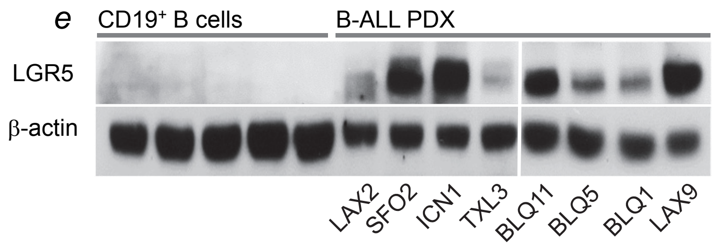


(**d**) mRNA levels of *LGR5* (top) and the PKA-scaffold *AKAP12* (bottom) in normal bone marrow (BM) cells were compared to *LGR5* and *AKAP12* mRNA levels in six cytogenetic subsets of B-ALL (*MLL*-rearranged, *BCR-ABL1*, hyperdiploid, *ETV6-RUNX1*, *TCF3-PBX1*), Burkitt's lymphoma, chronic lymphocytic leukemia (CLL), as well as T-ALL, myelodysplastic syndrome (MDS), chronic myeloid leukemia (CML) and acute myeloid leukemia (AML). B-ALL subtypes are highlighted in red.

(**e**) Western blot analyses of LGR5 protein levels in B-ALL xenografts (labels below) in comparison to normal CD19$^{+}$ B cells from peripheral blood are shown.

**Extended data figure 2:** *Identification of LGR5 as essential requirement for B-lymphopoiesis*

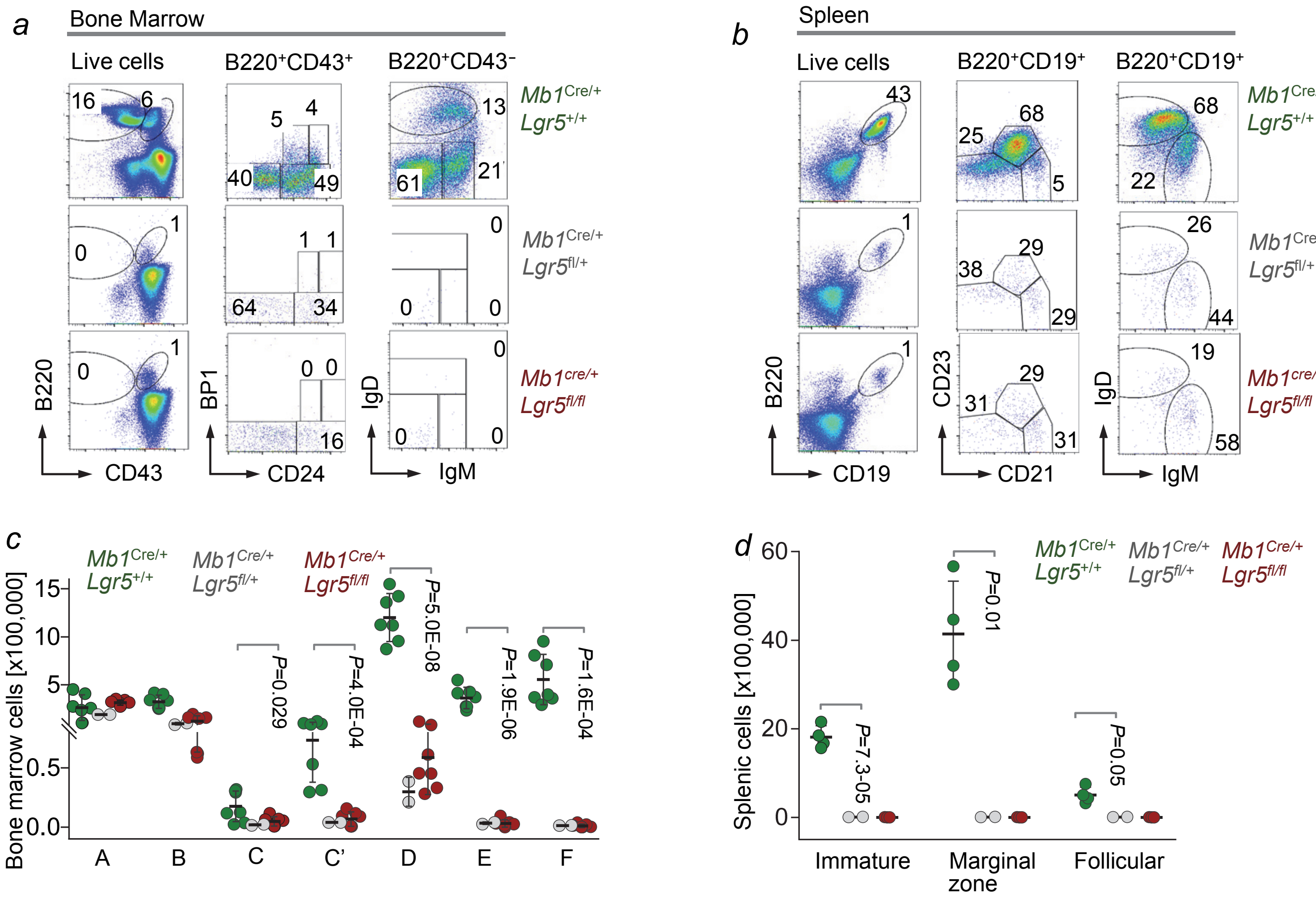


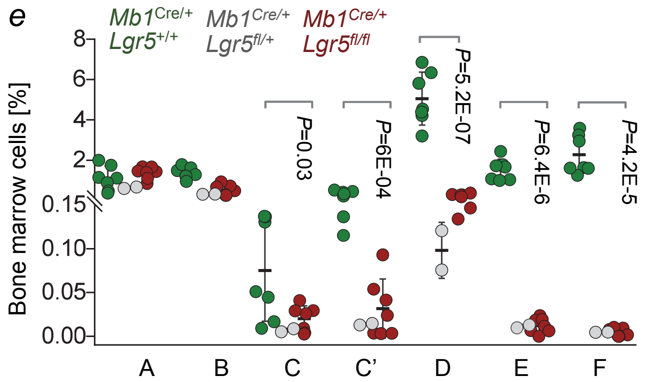


**a**) FACS analyses of Hardy Fractions (A-F) in the bone marrow of *Mb1*$^{Cre/+}$ *Lgr5*$^{+/+}$ (n=7), *Mb1*$^{Cre/+}$ *Lgr5*$^{fl/+}$ (n=2) and *Mb1*$^{Cre/+}$ *Lgr5*$^{fl/fl}$ (n=7) mice. B220$^{+}$ CD43$^{+}$ cells were further characterized as Fraction A (Bp1$^{-}$ CD24$^{-}$), B (Bp1$^{-}$ CD24$^{+}$), C (Bp1$^{lo}$ CD24$^{-}$), C' (Bp1$^{hi}$ CD24$^{-}$) and B220$^{+}$ CD43$^{-}$ cells were further divided into Fraction D (IgM$^{-}$ IgD$^{-}$), E (IgM$^{+}$ IgD$^{-}$) and F (IgM$^{+}$ IgD$^{+}$).

(**b**) Representative FACS analyses of splenic B-cell populations from *Mb1*$^{Cre/+}$ *Lgr5*$^{+/+}$ (n=4), *Mb1*$^{Cre/+}$ *Lgr5*$^{fl/+}$ (n=2) and *Mb1*$^{Cre/+}$ *Lgr5*$^{fl/fl}$ (n=3) mice. B220$^{+}$ CD19$^{+}$ B cells were further delineated as immature B cells (CD21$^{-}$ CD23$^{-}$), marginal zone B cells (CD21$^{+}$ CD23$^{-}$) and follicular B cells (CD21$^{+}$ CD23$^{+}$).

Absolute numbers of B-cell populations in the (**c**) bone marrow (**d**) spleens of *Mb1*$^{Cre/+}$ *Lgr5*$^{+/+}$ (green), *Mb1*$^{Cre/+}$ *Lgr5*$^{fl/+}$ (gray) and *Mb1*$^{Cre/+}$ *Lgr5*$^{fl/fl}$ (red) mice. (**e**) Frequencies of B cell developmental stages (Hardy Fractions A-F) within the bone marrow of above indicated mice.

**Extended data figure 3:** *Genetic deletion of Lgr5 does not affect mature B cells*

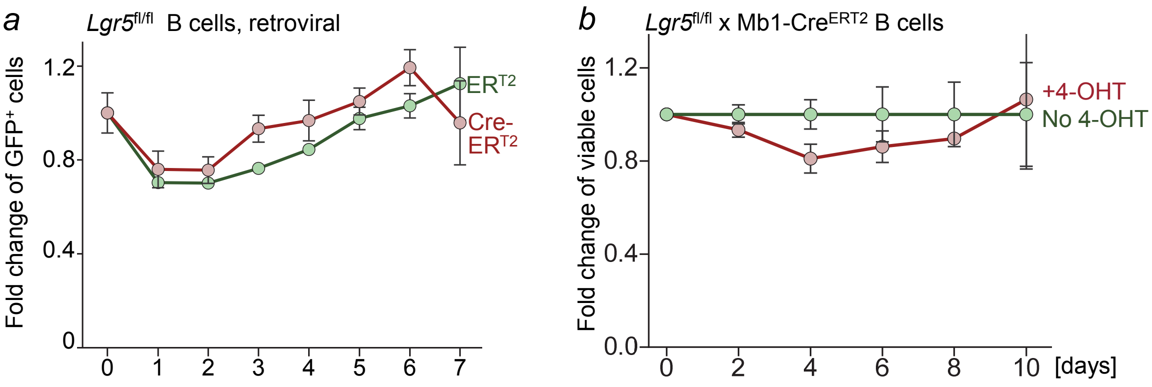


(**a**) Negatively selected B-cells from spleens of *Lgr5*$^{fl/fl}$ mice were transduced with GFP-tagged Cre-ER$^{T2}$ or ER$^{T2}$. Ratios of GFP$^{+}$ cells were monitored by FACS for 7 days after 4-OHT treatment.

(**b**) Splenic B cells from *Mb1*$^{CreERT2}$ *Lgr5*$^{fl/fl}$ mice were treated with or without 4-OHT and the number of viable cells were counted for 10 days.

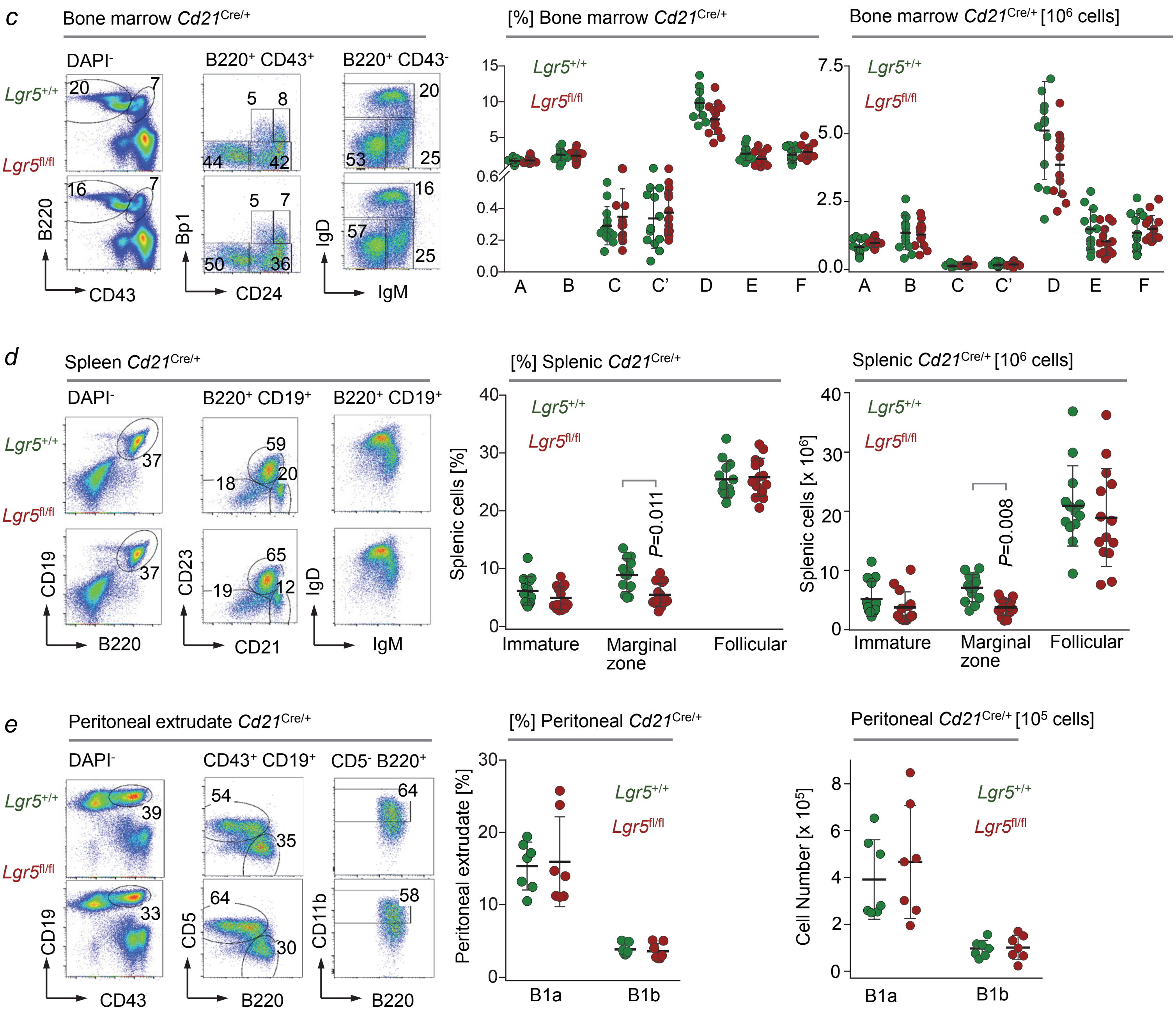


(**c**) Representative FACS analyses of B-cell development in the bone marrow of *Cd21*$^{Cre/+}$ *Lgr5*$^{+/+}$ (n=13) or *Cd21*$^{Cre/+}$ *Lgr5*$^{fl/fl}$ (n=14) mice was assessed by Hardy fraction stainings. Frequencies (middle) and absolute numbers (right) of B-cells in the bone marrow of *Cd21*$^{Cre/+}$ *Lgr5*$^{+/+}$ (green) or *Cd21*$^{Cre/+}$ *Lgr5*$^{fl/fl}$ (red) mice. (**d**) FACS analyses of B-cell populations in the spleens of *Cd21*$^{Cre/+}$ *Lgr5*$^{+/+}$ (green) or *Cd21*$^{Cre/+}$ *Lgr5*$^{fl/fl}$ (red) mice. Frequencies (middle) and absolute numbers (right) of splenic B-cells are shown.

(**e**) FACS analyses of B-cells in the peritoneal cavity of *Cd21*$^{Cre/+}$ *Lgr5*$^{+/+}$ (green; n=7) or *Cd21*$^{Cre/+}$ *Lgr5*$^{fl/fl}$ (red; n=7) mice. Frequencies (middle) and absolute numbers (right) of peritoneal cavity B1a (CD19$^{+}$ CD43$^{+}$ B220$^{+}$ CD5$^{+}$) and B1b cells (CD19$^{+}$ CD43$^{+}$ B220$^{+}$ CD5$^{-}$ CD11b$^{+}$) are detailed.

**Extended data figure 4:** *Lgr5 is not required for germinal center B-cell responses*

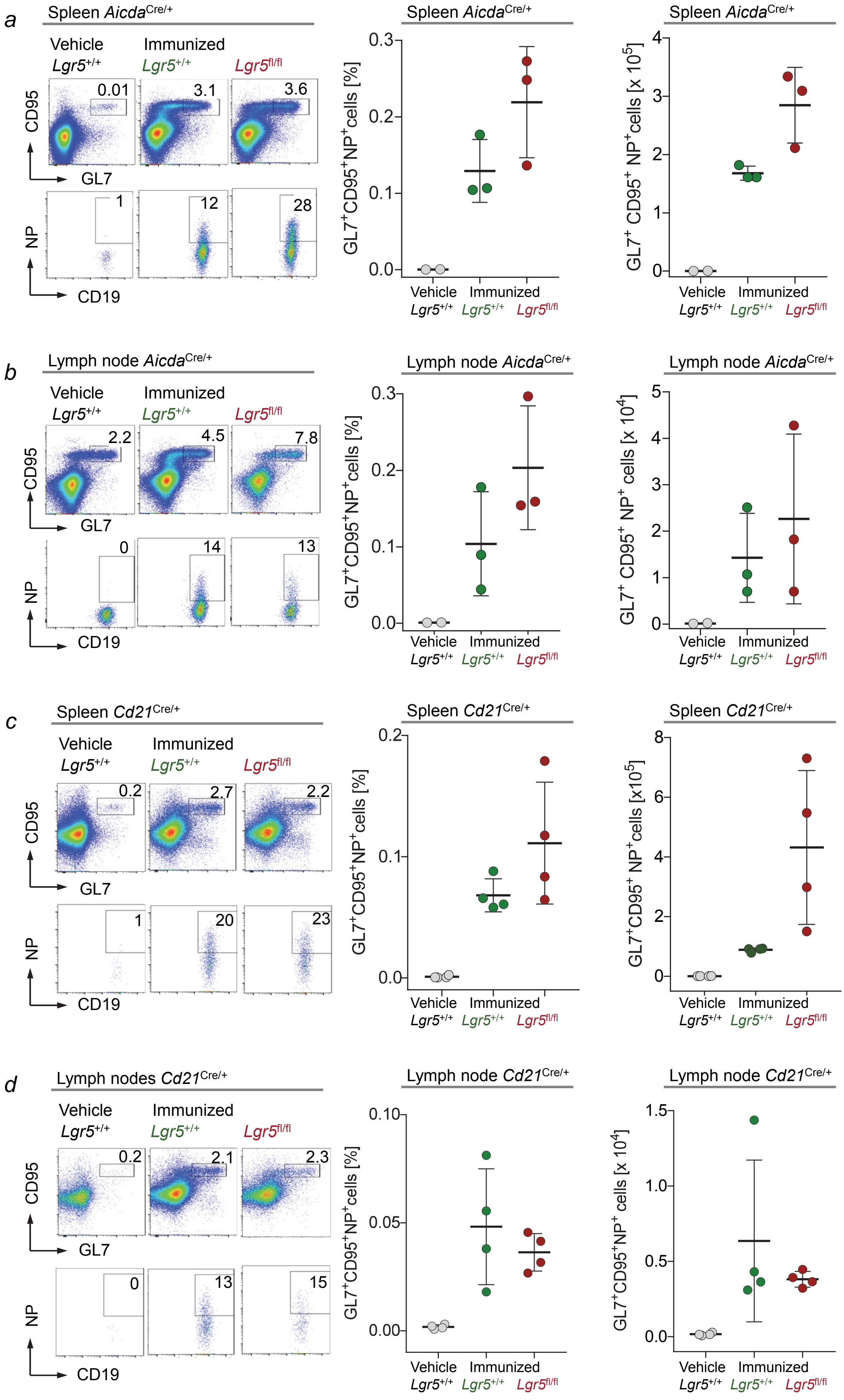


(**a**) FACS analyses of splenic B cells (gated on CD19$^+$ cells) from *Aicda*$^{Cre/+}$ *Lgr5*$^{+/+}$ (n=3, green) or *Aicda*$^{Cre/+}$ *Lgr5*$^{fl/fl}$ (n=3, red) mice that were injected with NP-hapten (days 7 and 12). *Aicda*$^{Cre/+}$ *Lgr5*$^{+/+}$ (n=2, gray) mice injected with vehicle were used as controls. Germinal center B-cells (CD19$^+$ CD95$^+$ GL7$^+$) were stained with fluorescent-labelled NP to identify antigen specific B cells. Frequencies (middle) and absolute numbers (right) of NP-specific germinal center B-cells in the spleen.

(**b**) FACS analyses of lymph node B cells (gated on CD19$^+$ cells) from immunized or non-immunized *Aicda*$^{Cre/+}$ *Lgr5*$^{+/+}$ *or Aicda*$^{Cre/+}$ *Lgr5*$^{fl/fl}$ mice. Frequencies (middle) and absolute numbers (right) of NP specific germinal center B cells in the lymph nodes.

(**c**) Flow cytometry analyses of spleens (gated on CD19$^+$ B cells) of *Cd21*$^{Cre/+}$ *Lgr5*$^{+/+}$ (n=4, green) *or* *Cd21*$^{Cre/+}$ *Lgr5*$^{fl/fl}$ (n=4, red) mice immunized with NP. *Cd21*$^{Cre/+}$ *Lgr5*$^{+/+}$ mice (n=4, gray) injected with vehicle were used as controls. Frequencies (middle) and absolute numbers (right) of NP-reactive germinal center B-cells in the spleens are shown.

(**d**) FACS analyses of lymph nodes (gated on CD19$^+$ B cells) from NP-immunized or control *Cd21*$^{Cre/+}$ *Lgr5*$^{+/+}$ *or* *Cd21*$^{cre/+}$ *Lgr5*$^{fl/fl}$ mice. Frequencies (middle) and absolute number (right) of NP-specific B-cells in lymph nodes from immunized and non-immunized mice (gray) are shown.

**Extended data figure 5:** *LGR5 mRNA levels predict poor clinical outcomes in B-ALL but not in mature B-cell malignancies and PTCL*

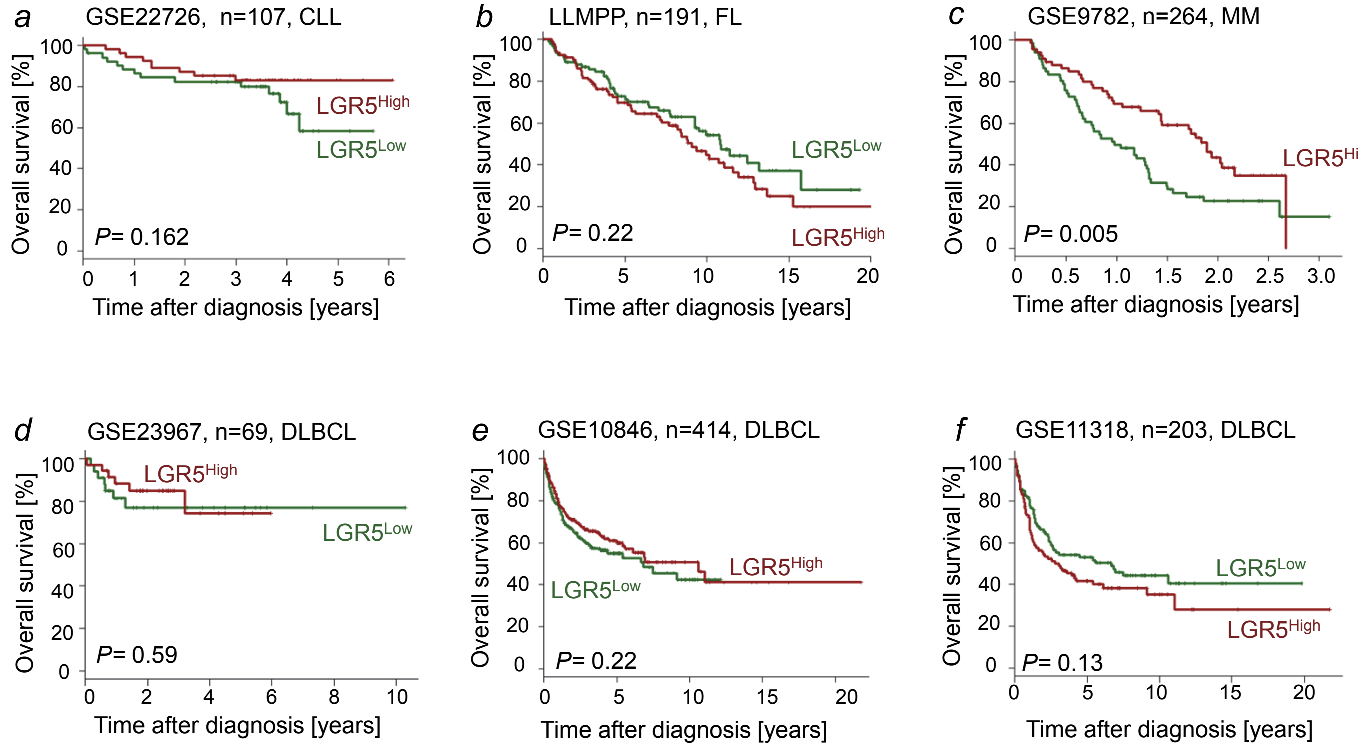


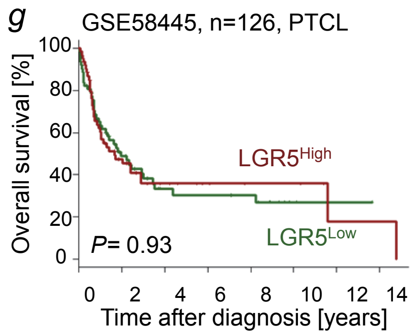


(**a-g**) Patients were divided into two groups based on higher (*LGR5*High) or lower (*LGR5*Low) than median *LGR5* mRNA levels. Overall survival (OS) was assessed in the two groups by Kaplan-Meier analysis and *P*-values were calculated by log-rank test in clinical cohorts of patients with (**a**) Chronic lymphocytic leukemia (CLL, n=107, *P*=0.162) (**b**) Follicular lymphoma (FL, n=191, *P*=0.22), (**c**) Multiple myeloma (n=264, *P*=0.005), and three clinical cohorts of patients with Diffuse large B cell lymphoma, (**d**) (DLBCL, n=69, *P*=0.59), (**e**) DLBCL (n=414, *P*=0.22), (**f**) DLBCL(n=203, *P*=0.13), (**g**) peripheral T cell lymphoma (PTCL, n=126, *P*=0.93)

**Extended data figure 6:** *Lgr5-deletion reduces competitive fitness of B-ALL cells and prolongs latency in transplant recipients*

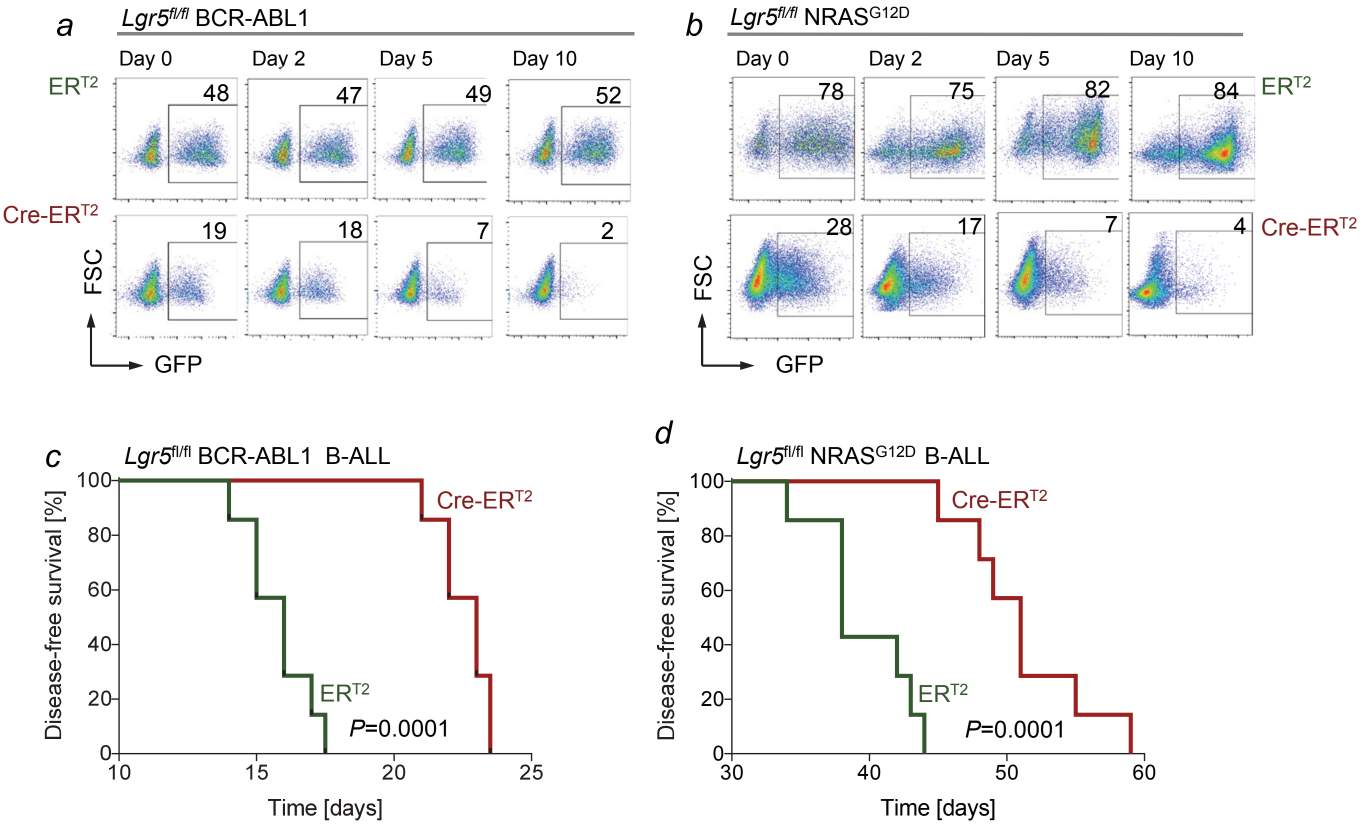


(**a**) *BCR-ABL1* or (**b**) *NRAS*$^{G12D}$-transformed *Lgr5*$^{fl/fl}$ pre-B cells gave rise to B-ALL, were transduced with GFP-tagged, 4-hydroxy-tamoxifen (4-OHT)-inducible Cre-ER$^{T2}$ or ER$^{T2}$ empty vector. Representative FACS analysis to assess competitive fitness or depletion of GFP$^{+}$ cells (n=3). (**c-d**) Sublethally irradiated (2 Gy) NSG mice were transplanted with (**c**) 300,000 *BCR-ABL1* (**d**) 1 million *NRAS*$^{G12D}$-transformed *Lgr5*$^{fl/fl}$ B-ALL expressing Cre-ER$^{T2}$ or ER$^{T2}$. Kaplan-Meier analysis was performed to compare survival for recipient NSG mice (n=7, *P*=0.0001, log-rank test).

**Extended data figure 7:** *LGR5 promotes self-renewal and leukemia-initiation in patient-derived B-ALL cells*

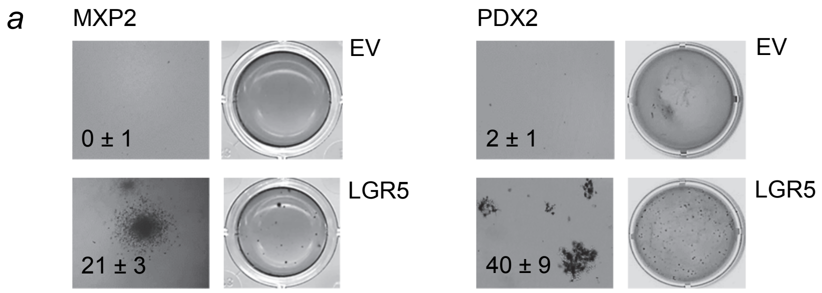


(**a**) Patient-derived B-ALL cells (MXP2, PDX2) were transduced with LGR5 or empty vector controls (EV) and plated on methylcellulose, colonies were counted 14 days later. (**b**) Patient-derived B-ALL cells were transduced with firefly luciferase and transduced with ds-Red *LGR5* or EV. ds-Red$^{+}$ cells were sorted by FACS and transplanted into sublethally irradiated (2 Gy) NSG recipients. (**c**) Kaplan-Meier analyses show overall survival for each group (n=7) of recipient mice.

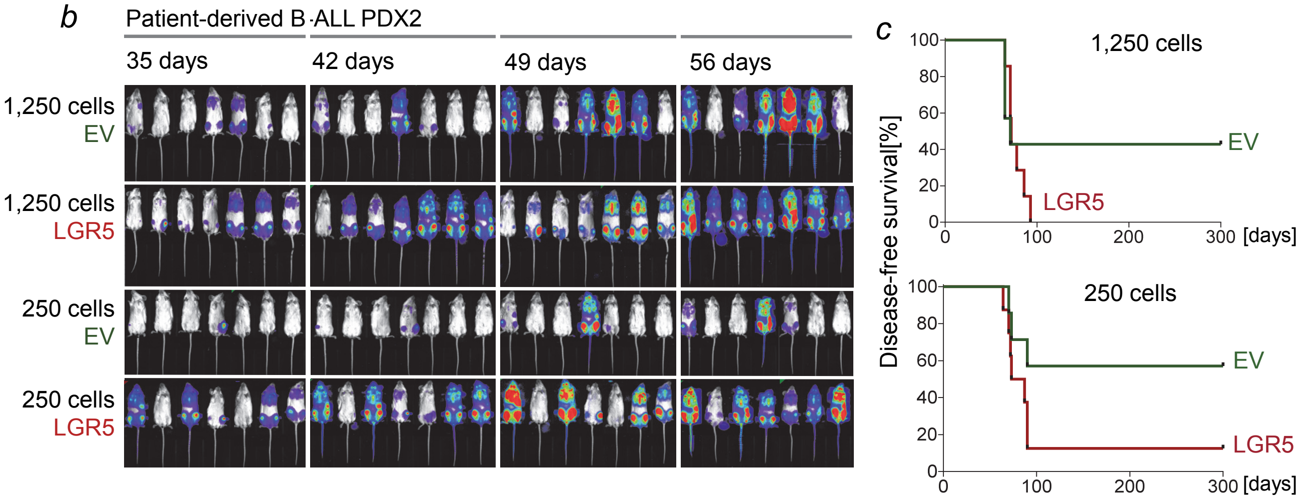


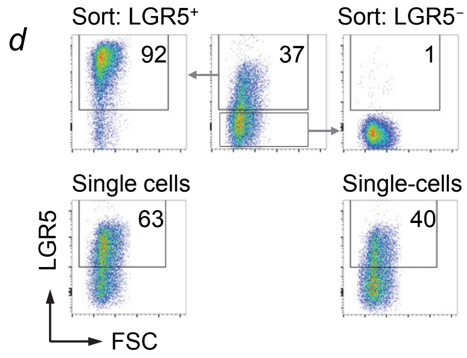


(**d**) Patient-derived B-ALL cells were sorted based on surface LGR5 expression as single LGR5$^{+}$ and LGR5$^{-}$ cells. After two weeks, outgrowth from single sorted cells was observed and FACS analysis was repeated to determine LGR5 surface expression levels.

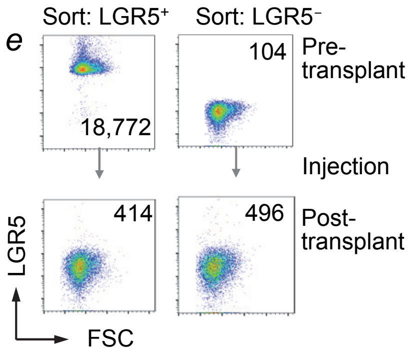


(**e**) Cell sorting of LGR5$^{+}$ and LGR5$^{-}$ B-ALL cells, and flow cytometry of LGR5 surface levels before and after injection into NSG mice.

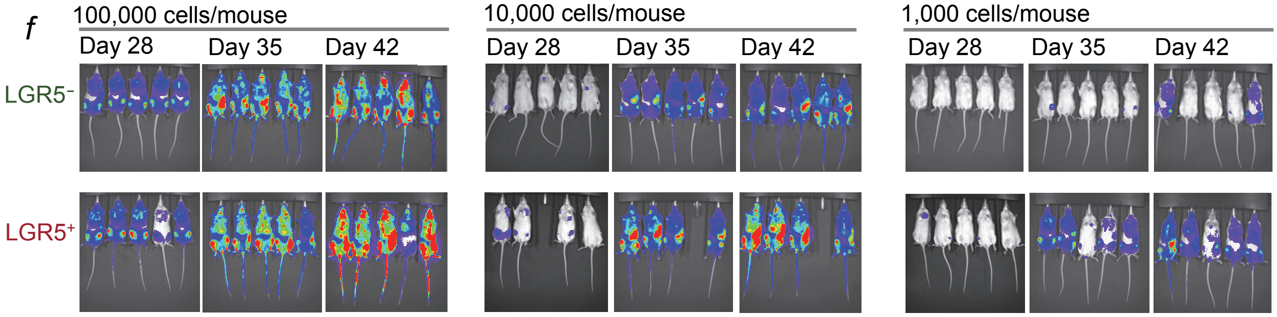


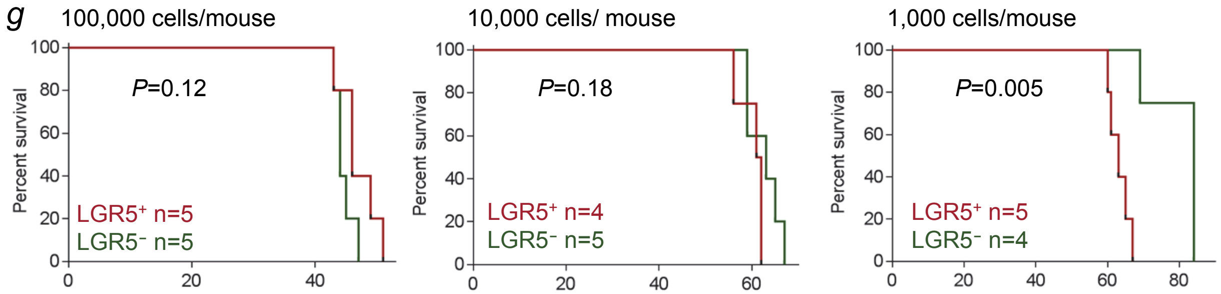


Firefly luciferase-labeled B-ALL PDX cells were sorted based on LGR5 expression (**d**, **e**). $10^3$, $10^4$ or $10^5$ LGR5$^{+}$ and LGR5$^{-}$ cells were injected into sub-lethally irradiated (2 Gy) NSG mice.

(**f**) Leukemia burden was assessed by bioluminescence imaging 28, 35 and 42 days after injection. (**g**) Kaplan-Meier analyses show overall survival of transplant recipient mice (n=5) transplanted with $10^3$, $10^4$ or $10^5$ LGR5$^{+}$ and LGR5$^{-}$ B-ALL cells (x-axis, days after injection).

**Extended data figure 8:** *Bio-ID analyses to identify LGR5-interaction partners in human B-ALL cells*

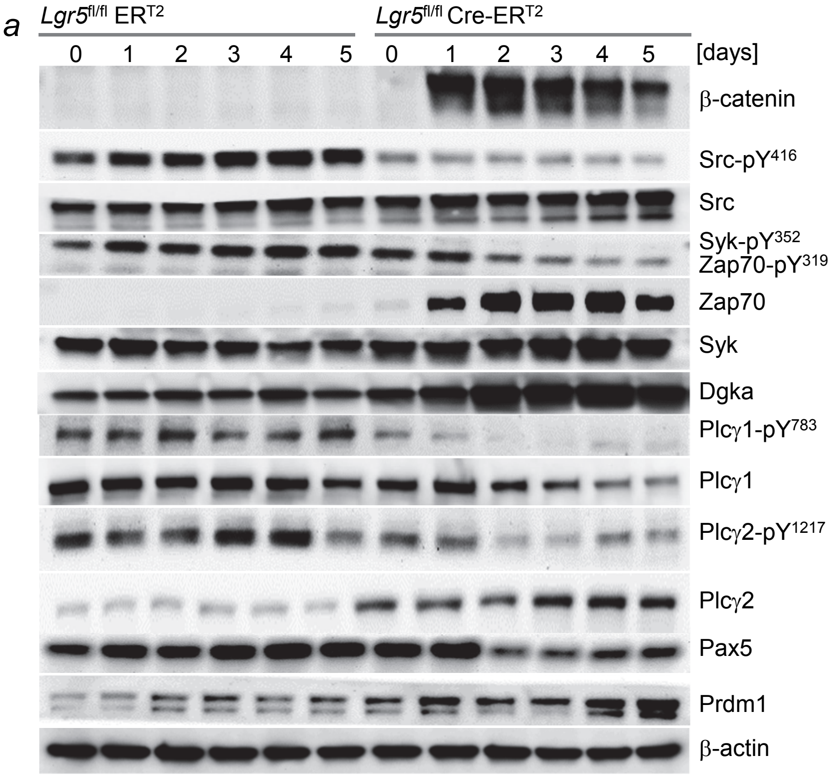


(**a**) To determine how LGR5-deletion affected typical signaling pathways in B-ALL cells, Western blot analyses were performed in *Lgr5*$^{fl/fl}$ *BCR-ABL1* B-ALL cells expressing ER$^{T2}$ or Cre-ER$^{T2}$, in a time course analysis (0 to 5 days) following 4-OHT addition. Protein levels of β-catenin, Src-pY$^{416}$, global Src, Syk-pY$^{352}$/ Zap70-pY$^{319}$, global Zap70, global Syk, Dgkα, Plcγ1-pY$^{783}$, global Plcγ1, Plcγ2-pY$^{1217}$, global Plcγ2, Pax5 and Prdm1 were studied.

These studies with a focus on tyrosine phosphorylation, were performed because our phospho-proteomic analysis preferentially detected S- and T-phosphorylation events. S/T-phosphorylation occurs at greater abundance than tyrosine-phosphorylation.

(**b, c**) SUPB15 (right) and PDX2 (left) cells were transduced with ds-Red labeled LGR5 fused with its cytoplasmic C-terminus to HA-tag and the biotin-ligase BirA. Cells transduced with EV (HA-tag) and BirA were used as control. FACS-sorted ds-Red$^{+}$ cells were incubated overnight with 50 µmol l$^{-1}$ biotin.

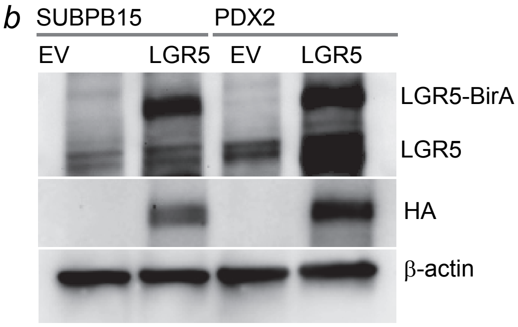


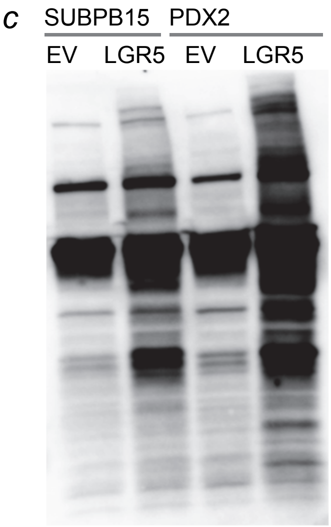


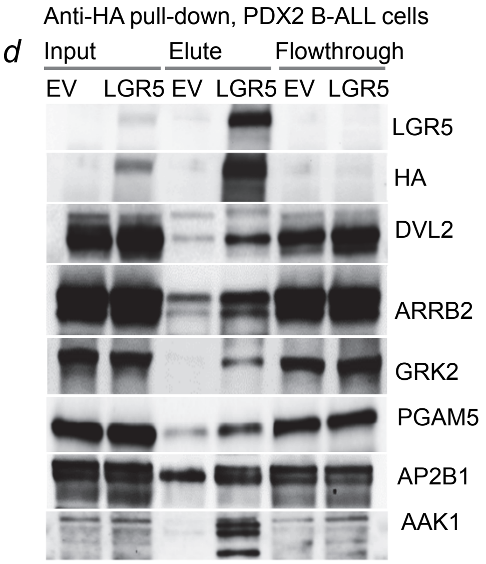


(**b, c**) blot analyses were performed to confirm over-expression of (**b**) LGR5 and HA-tag and (**c**) Streptavidin-HRP (SA-HRP) was used to asses biotinylation of proteins by LGR5-BirA. (**d**) Pull down experiment was performed with PDX2 cell expressing HA-tagged LGR5 or EV (input).
Western blot analyses were performed to identify proteins bound to (elute) and not bound (flow through) anti-HA beads.

(**e-f**) Scenario of LGR5-mediated negative regulation of WNT/β-catenin signaling in pre-B cells and B-ALL: FZD and LGR5 are structurally similar GPCRs and compete for binding to the proximal scaffold disheveled2 (DVL2).

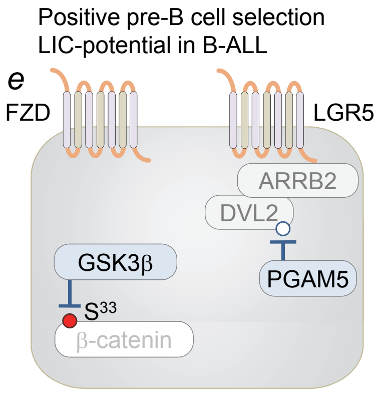


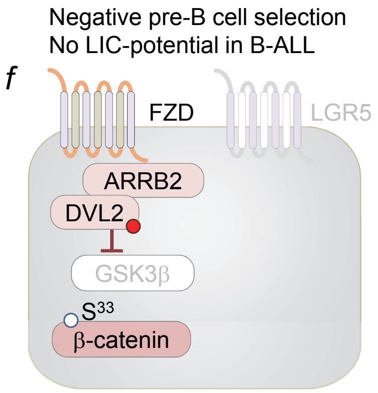


In positively selected pre-B cells that express a functional pre-BCR and B-ALL cells expressing an oncogenic pre-BCR mimic, high expression levels of LGR5 lead to sequestration of DVL2 from FZD receptors (**e**).

In proximity to LGR5, DVL2 is dephosphorylated by PGAM5 (**e**), which disrupts the ability of DVL2 to inhibit GSK3β, which then phosphorylates and destabilizes β-catenin. (**f**) Upon *Lgr5*-deletion, DVL2 is released and associates with FZD receptors is constitutively phosphorylated, inhibits GSK3β, which results in accumulation of β-catenin.

**Extended data figure 9:** *Genetic hyperactivation of β-catenin represents the functional equivalent of Lgr5-deletion in pre-B cells*

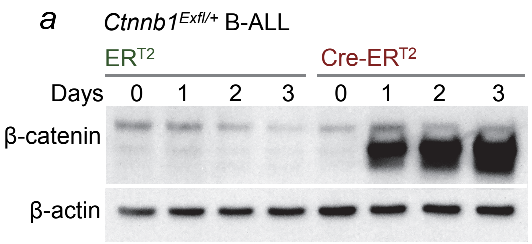


*BCR-ABL1*-driven *Ctnnb1*$^{ex3fl/+}$ B-ALL cells and *Lgr5*$^{fl/fl}$ B-ALL cells were transduced with GFP-tagged 4-OHT-inducible Cre-ER$^{T2}$ or ER$^{T2}$. (**a**) Western blot analyses were performed to quantify β-catenin levels upon 4-OHT treatment in *Ctnnb1*$^{Ex3fl/+}$ B-ALL cells. (**b-c**) Representative FACS analysis to measure the frequencies of GFP$^{+}$ cells following 4-OHT induction in (**b**) *Lgr5*$^{fl/fl}$ B-ALL and (**c**) *Ctnnb1*$^{Ex3fl/+}$ B-ALL cells.

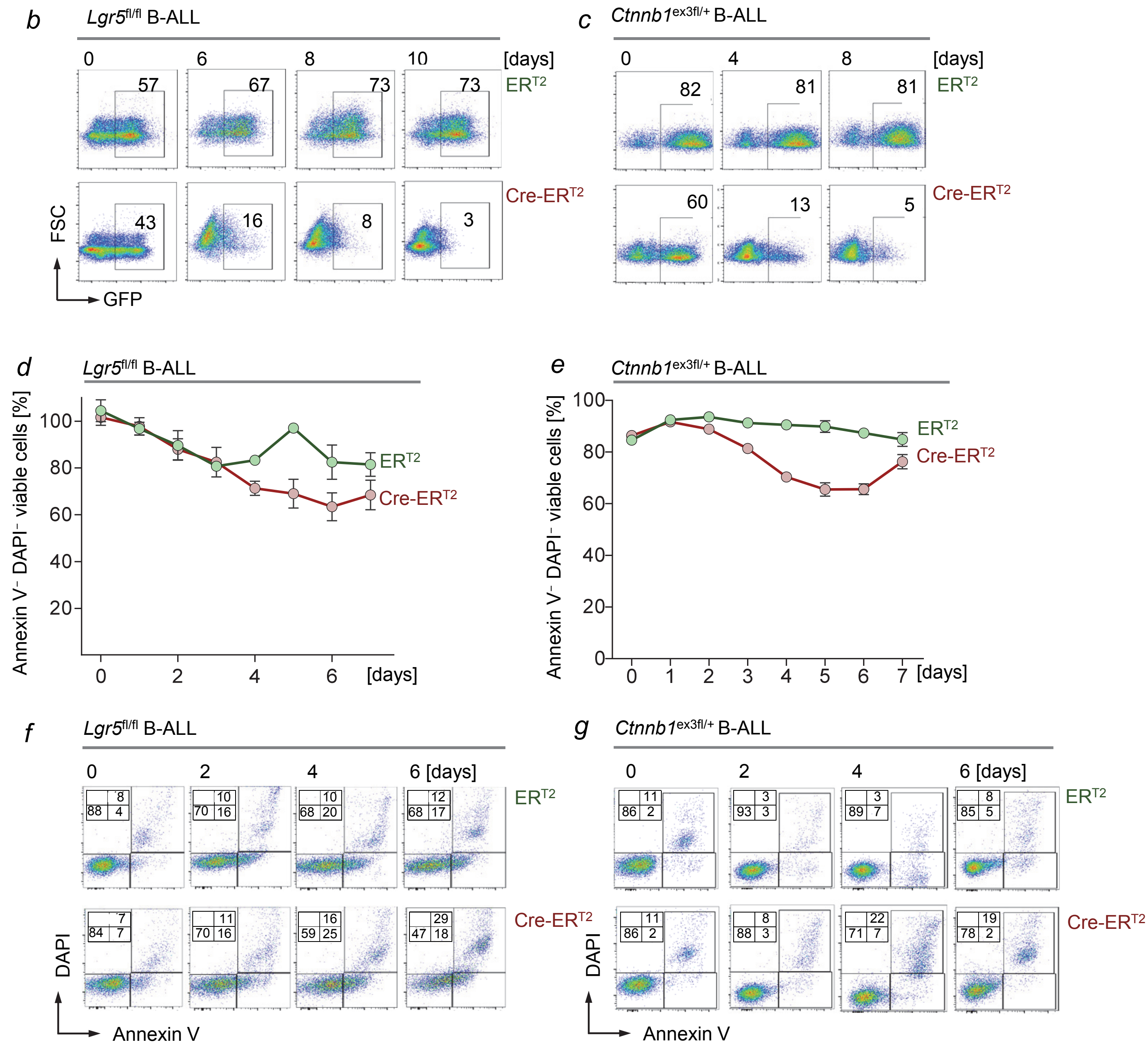


Annexin V and DAPI stainings were performed for *Lgr5*$^{fl/fl}$ B-ALL cells and *Ctnnb1*$^{ex3fl/+}$ B-ALL cells carrying Cre-ER$^{T2}$ or ERT2 empty vectors following 0 to 7 days after 4-OHT addition. **(d-e)** Frequencies of viable cells (DAPI$^{-}$ Annexin V$^{-}$) were measured daily for the times indicated after 4-OHT induction. **(f-g)** Representative FACS analysis for Annexin V and DAPI stainings are shown.

**Extended data figure 10:** *β-catenin deletion rescues loss of Lgr5 in B-ALL cells*

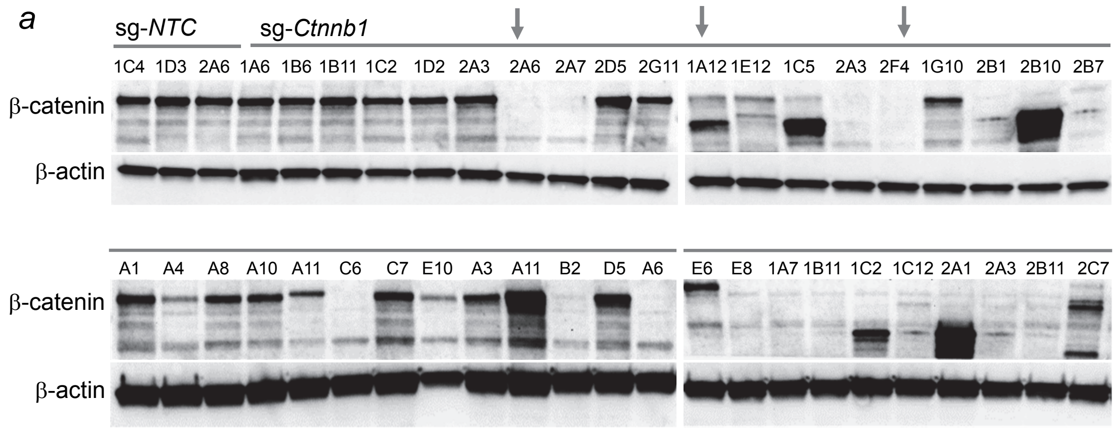


(**a**) *BCR-ABL1* transformed *Lgr5*$^{fl/fl}$ B-ALL cells were transduced with Cas9 and guide RNAs against *Ctnnb1* (sg-*Ctnnb1*) or non-targeting control (sg-NTC). Single cell-derived colonies were grown from these cells and Western blot analyses were performed to identify colonies with deletion of β-catenin.

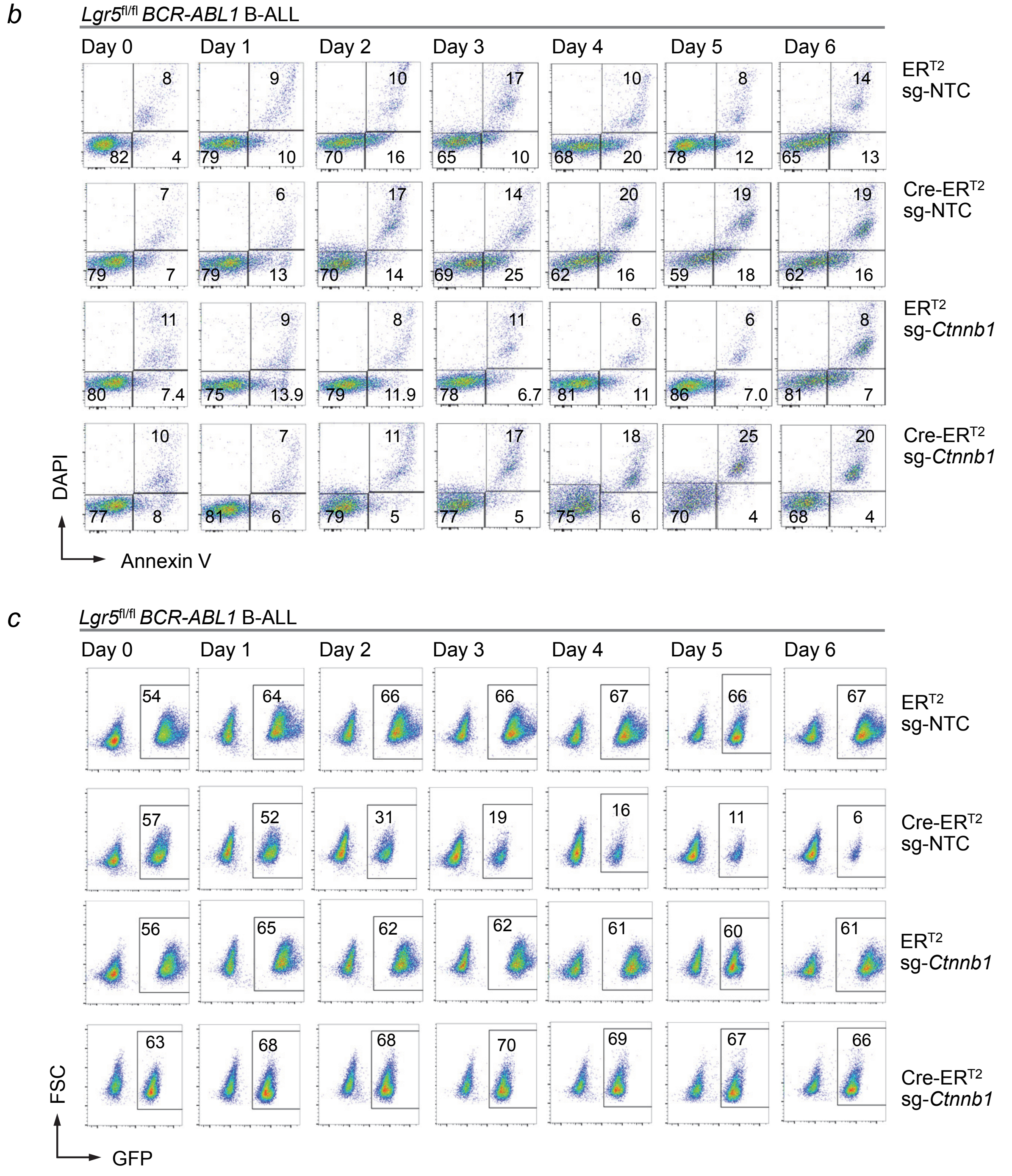


(**b**) Representative FACS plots for Annexin V and 7AAD staining of *BCR-ABL1* transformed *Lgr5*$^{fl/fl}$ B-ALL cells with (sg-*Ctnnb1*) or without (sg-NTC) β-catenin deletion and carrying Cre-ER$^{T2}$ or ER$^{T2}$. (**C**) GFP$^{+}$ CRISPR-Cas9 edited cells were mixed with unedited and unlabeled cells and FACS analyses were performed to quantify the changes of the frequency of GFP$^{+}$ cells.

**Extended data figure 11:** *Lgr5 functions as a negative regulator of β-catenin in B-cells but not in colon carcinoma cells*

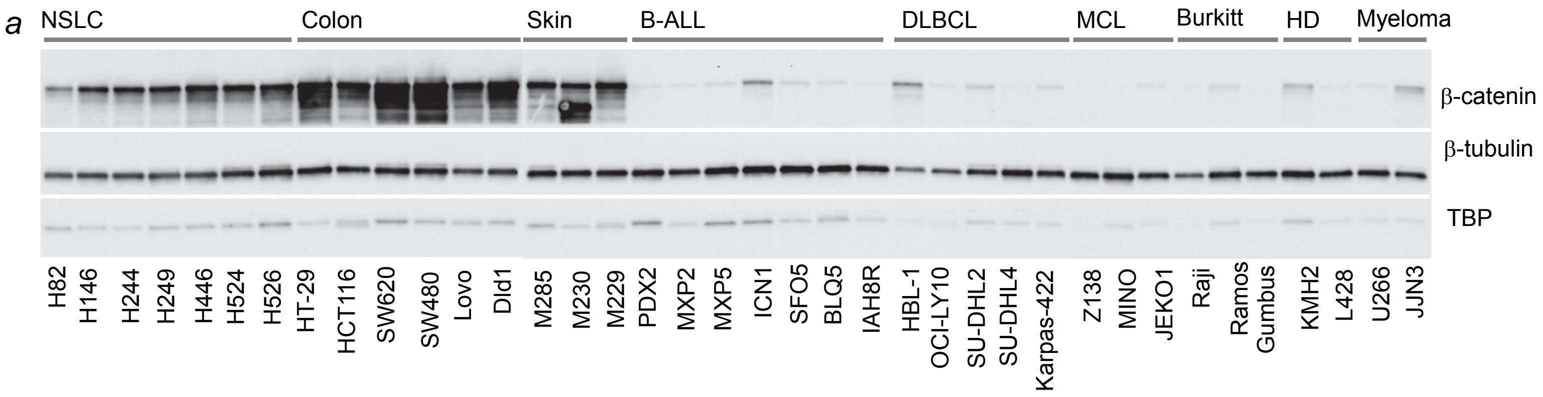


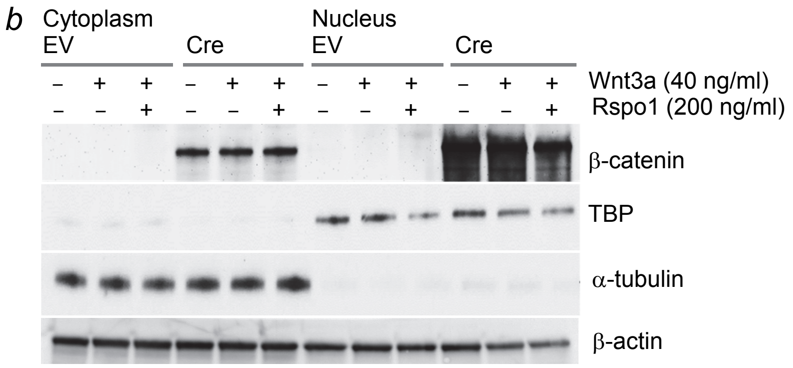


(**a**) Western blot analyses on nuclear fractions of non-small lung carcinoma (NSCLC), colon cancer, malignant melanoma leukemia and B-ALL, diffuse large B-cell lymphoma (DLBCL), mantle cell lymphoma (MCL), Burkitt's, Hodgkin's disease (HD) and multiple myeloma cell lines for β-catenin, β-tubulin and TBP. Western blots of the nuclear fractions from the same cell lysates are shown in **Fig. 5e**.

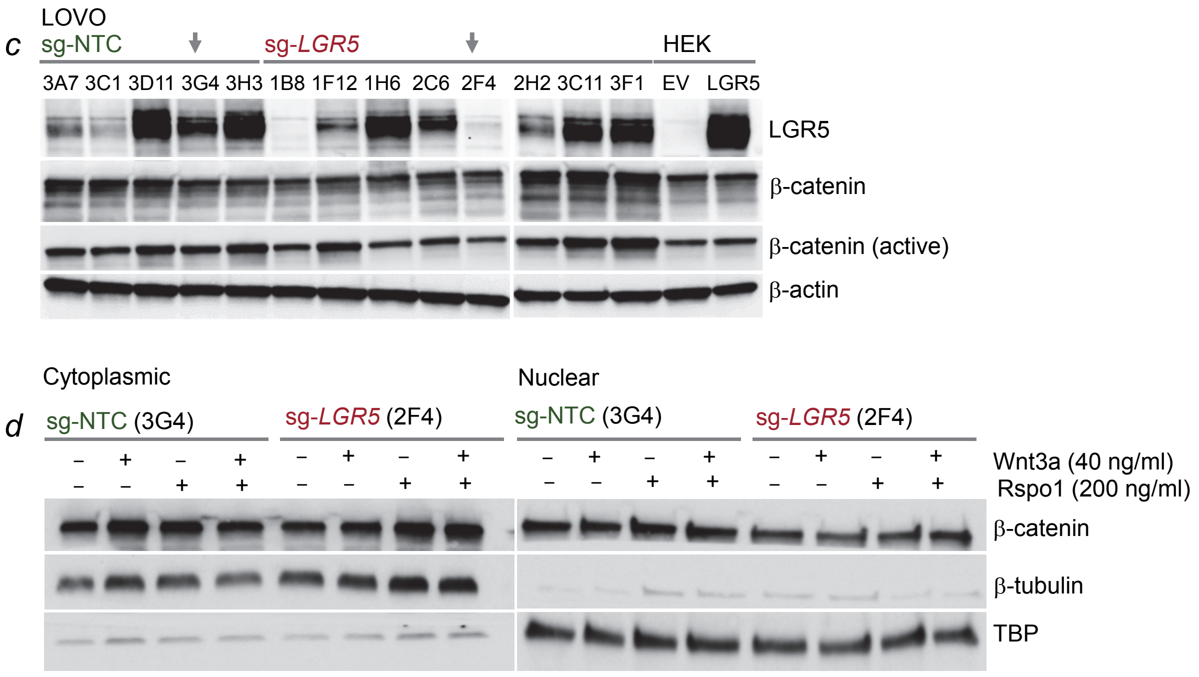


(**b**) *BCR-ABL1*-transformed *Lgr5*$^{fl/fl}$ pre-B ALL cells were treated with 40 ng ml$^{-1}$ Wnt3a and/or 200 ng ml$^{-1}$ R-Spondin-1 (Rspo1) for 16 hours and levels of β-catenin in the cytoplasm and nucleus were determined 4 days after 4-OHT induction. (**c**) Colon cancer cells (LOVO) were electroporated with Cas9 ribonucleoproteins (RNPs), complexes of recombinant Cas9 with non-targeting (sg-NTC) crRNAs or crRNAs targeting *LGR5* (sg-*LGR5*). Single cell-derived colonies were grown from Cas9-RNP electroporated cells and Western blot analyses were performed to measure the levels of LGR5, β-catenin, active (or stable) β-catenin (non-phosphorylated $S^{33}/S^{37}/T^{41}$) and β-actin as loading control. (**d**) Colon cancer cells (LOVO) grown from single colonies were treated with 40 ng ml$^{-1}$ Wnt3a and/or 200 ng ml$^{-1}$ Rspo1 for 16 hours and the levels of nuclear and cytoplasmic β-catenin were determined by Western blot analyses.

**Extended data figure 12:** *Deletion of catalytic subunit of PKA partially mitigates loss of Lgr5 in B-ALL cells*

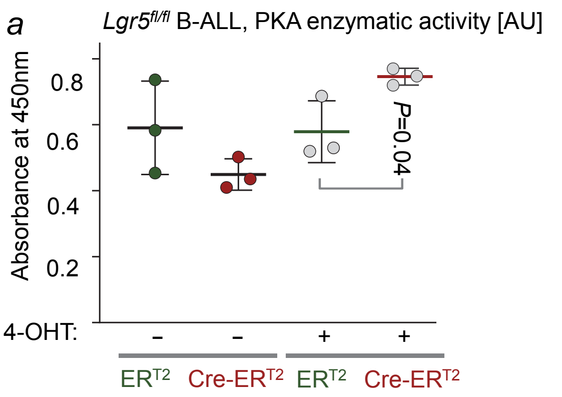


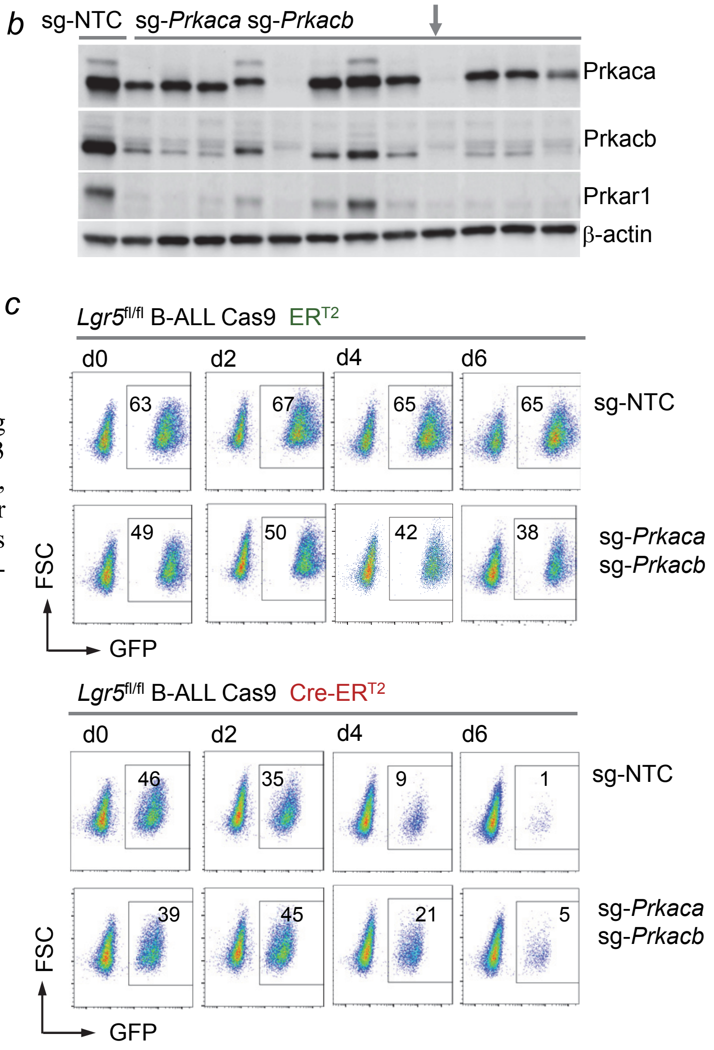


(**a**) *BCR-ABL1* transformed *Lgr5*$^{fl/fl}$ B-ALL cells expressing Cre-ER$^{T2}$ or ER$^{T2}$ were treated with 4-OHT or vehicle for 3 days and protein kinase A (PKA) activity was measured. AU, arbitrary units. (**b**) *Lgr5*$^{fl/fl}$ B-ALL cells expressing Cre-ER$^{T2}$ or ER$^{T2}$ were subsequently transduced with Cas9 and guide RNAs against *Prkaca* and *Prkacb* (sg*Prkaca*, sg*Prkacb*) or non-targeting controls (sgNTC).

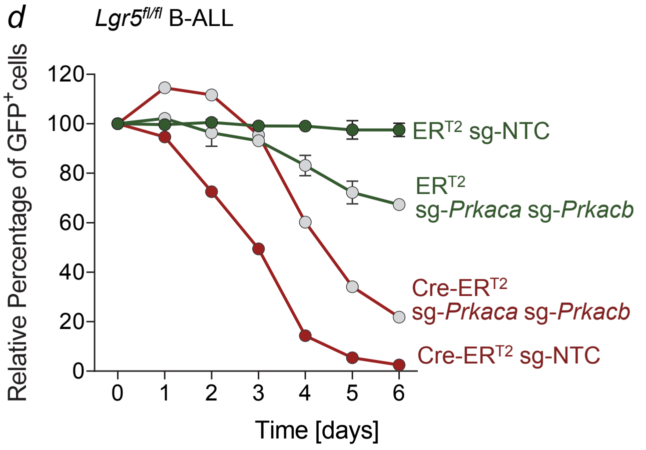


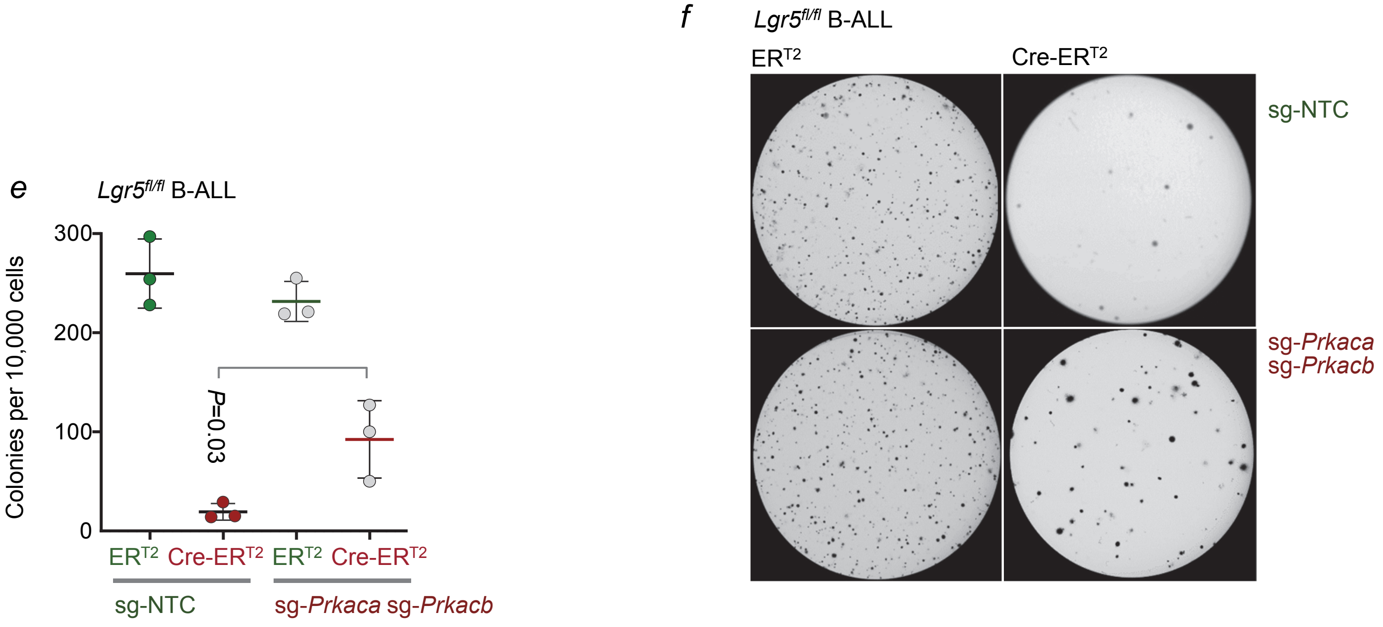


(**b**) Single cell-derived colonies were grown from these cells and Western blot analyses were performed to identify colonies with deletions of both catalytic subunits *Prkaca* and *Prkacb*. (**c-d**) GFP-tagged CRISPR-Cas9 edited cells with deletions of *Prkaca* and *Prkacb* were mixed with unedited and unlabeled B-ALL cells. FACS analyses were performed to quantify competitive fitness or depletion of GFP$^{+}$ cells. (**e-f**) *BCR-ABL1* transformed *Lgr5*$^{fl/fl}$ B-ALL cells expressing Cre-ER$^{T2}$ or ER$^{T2}$ that carry deletion of the catalytic PKA-subunits (sg-*Prkaca* sg*Prkacb*) or no deletion (sg-NTC) were plated on methylcellulose 2 days after 4-OHT induction. (**e**) Number of colonies per 10,000 cells is shown. (**f**) Representative images of colonies were taken 10 days after plating (n=3).

**Extended data figure 13:** *Deletion of Rhoa does not mitigate loss of Lgr5 in B-ALL cells*

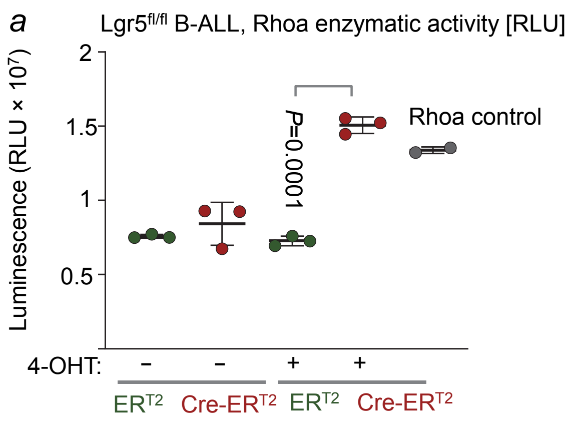


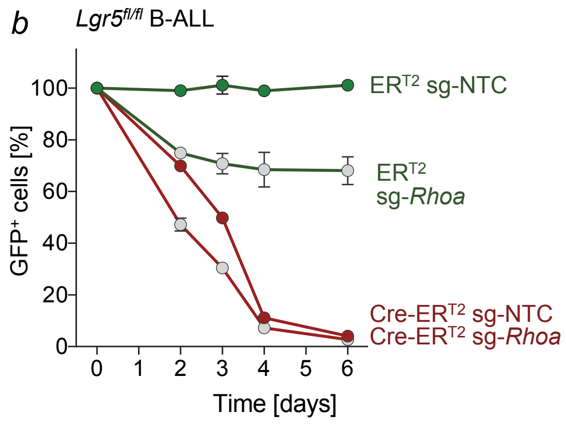


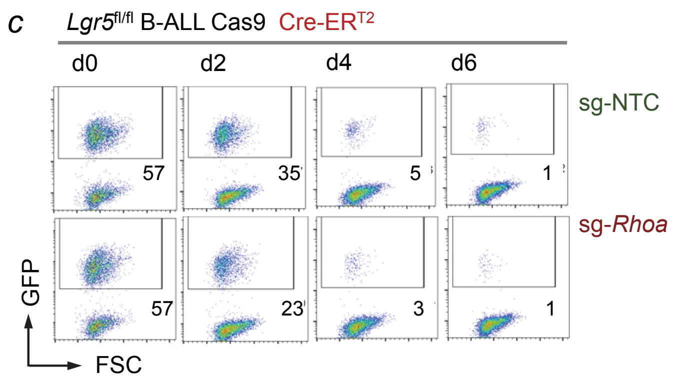


(**a**) *BCR-ABL1 Lgr5*$^{fl/fl}$ B-ALL cells expressing Cre-ER$^{T2}$ or ER$^{T2}$ were treated with 4-OHT or vehicle for 3 days and Rhoa-activity was measured.

(**b, c**) *BCR-ABL1* transformed *Lgr5*$^{fl/fl}$ B-ALL cells with Cre-ER$^{T2}$ or ER$^{T2}$ were edited with guide RNA against *Rhoa* (sg-*Rhoa*) or control guides (sg-NTC). GFP$^{+}$ cells with Cas9-RNP mediated deletion of *Rhoa* (sg-*Rhoa*) were mixed with unedited (sg-NTC) unlabeled cells and competitive fitness or depletion of GFP$^{+}$ cells was assessed by FACS for 6 days after 4-OHT treatment.

(**d**) 10,000 cells of *Lgr5*$^{fl/fl}$ B-ALL cells with Cre-ER$^{T2}$ or ER$^{T2}$, with (sg-*Rhoa*) and without (sg-NTC) *Rhoa*-deletion, were plated on semisolid methylcellulose agar 2 days after 4-OHT induction. Colonies were counted and imaged 10 days after plating (n=3). (**d**) Statistical analysis of colony numbers per 10,000 plated cells and representative images of colonies are shown.

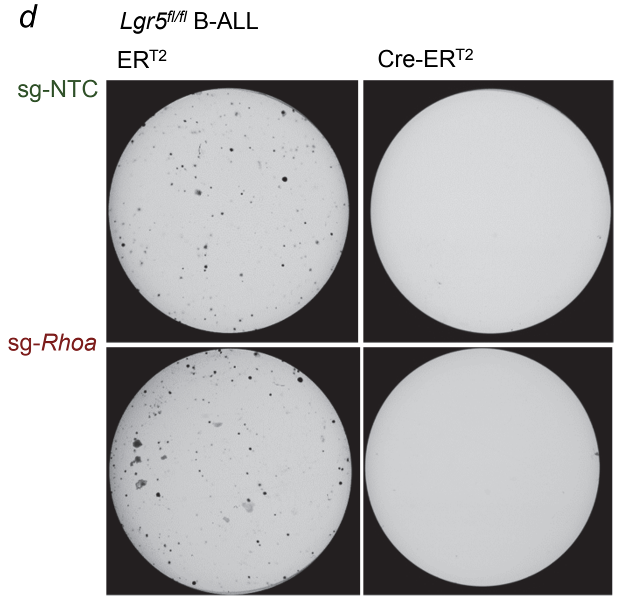


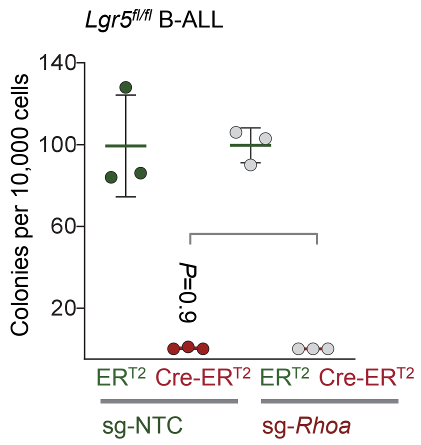

**Extended data figure 14:** *Overexpression of Myc rescues loss of Lgr5 and β-catenin hyperactivation in B-ALL cells*

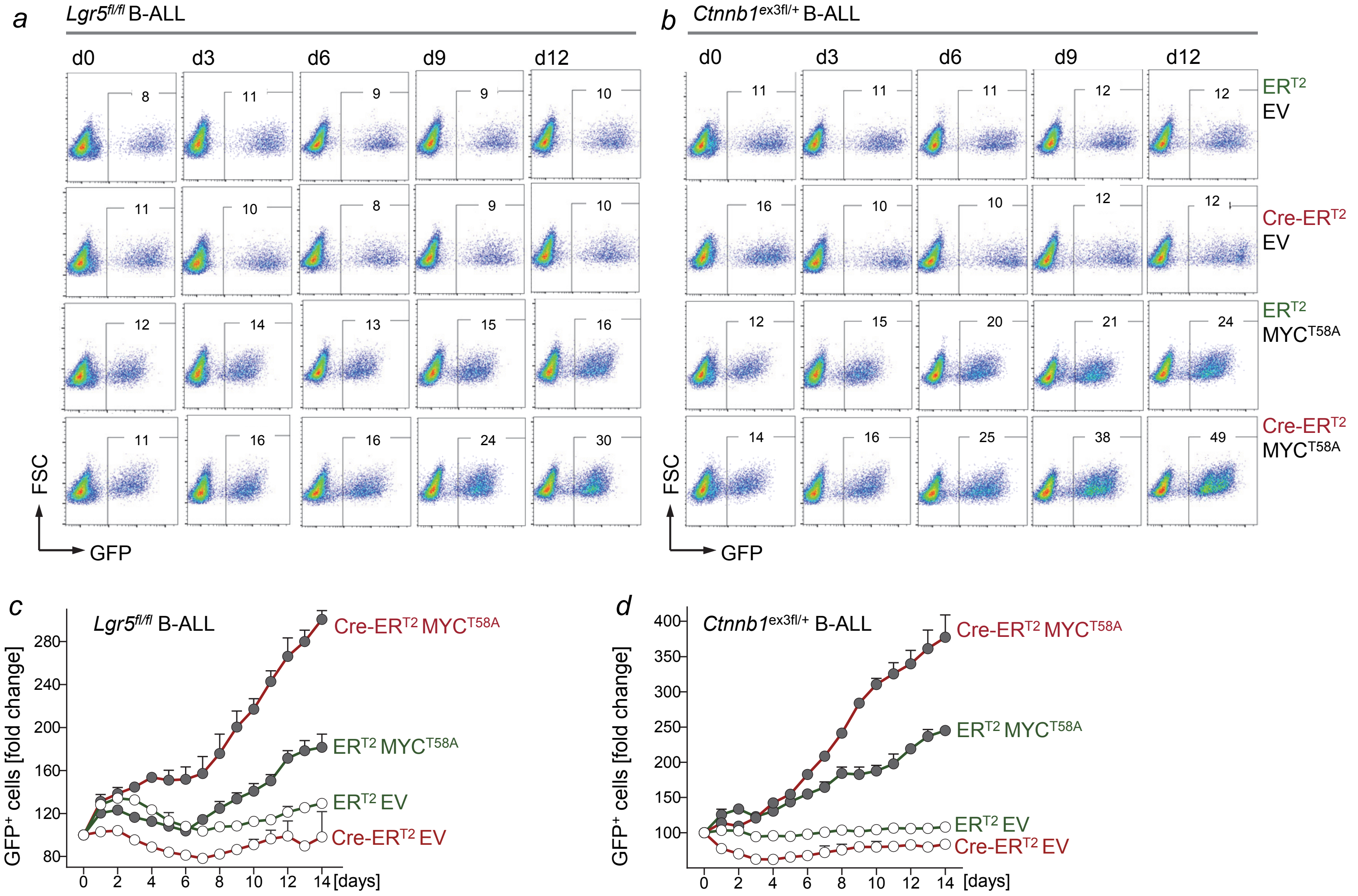


*BCR-ABL1* transformed $Lgr5^{fl/fl}$ (**a**) and $Ctnnb1^{ex3fl/+}$ (**b**) B-ALL cells expressing 4-OHT-inducible Cre-$ER^{T2}$ or $ER^{T2}$ were transduced with GFP-tagged degradation-resistant $Myc^{T58A}$ or EV. FACS analyses were performed to determine the enrichment or depletion of the $GFP^+$ cells upon 4-OHT mediated activation of Cre and FACS plots for days 0, 3, 6, 9 and 12 are shown for $Lgr5^{fl/fl}$ (**a**) and $Ctnnb1^{ex3fl/+}$ (**b**) B-ALL cells. Graphic representation of the time course analysis revealed that while Myc-overexpression increased competitive fitness of B-ALL cells, the competitive advantage was substantially widened upon *Lgr5*-deletion (**c**) or β-catenin hyperactivation (**d**).

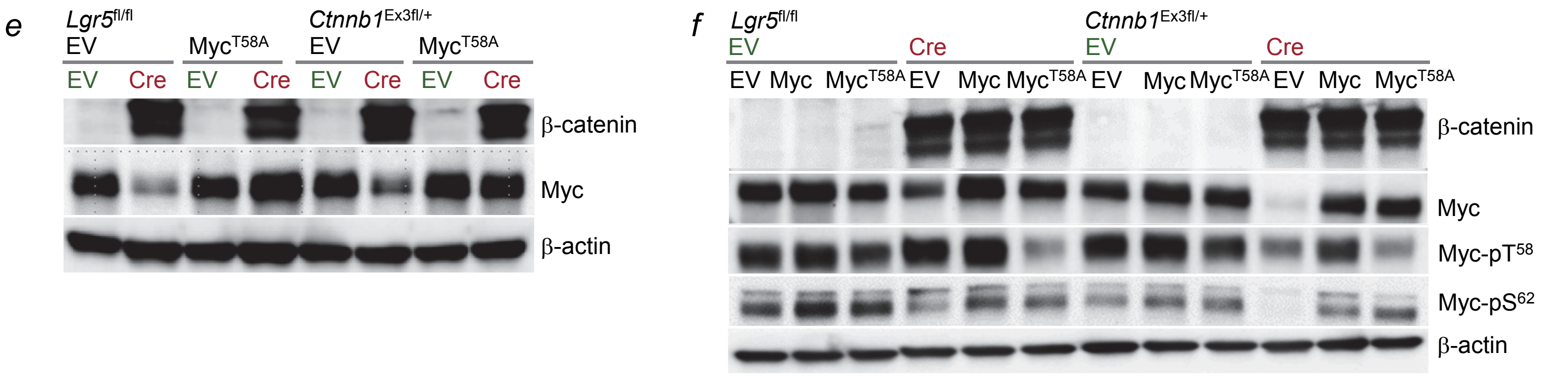


To confirm the identify of the cells studied in (**a-d**), $Lgr5^{fl/fl}$ and $Ctnnb1^{ex3fl/+}$ B-ALL cells transduced with GFP-tagged $Myc^{T58A}$ or EV were FACS-sorted for $GFP^+$ cells and Western blot analyses were performed for β-catenin and Myc levels 3 days after 4-OHT treatment (**e**). *BCR-ABL1* transformed $Lgr5^{fl/fl}$ and $Ctnnb1^{ex3fl/+}$ B-ALL cells expressing Cre-$ER^{T2}$ or $ER^{T2}$ were transduced with GFP-tagged Myc, degradation-resistant $Myc^{T58A}$ or EV. Western blot analyses for β-catenin, global Myc, Myc-$pT^{58}$, Myc-$pS^{62}$ were performed for FACS-sorted $GFP^+$ cells 3 days after 4-OHT treatment (**f**).

**Extended data figure 15:** *β-catenin associates with the pre-B cell factor Ikzf1 to promote an Ikzf1 transcriptional program, including repression of Myc*

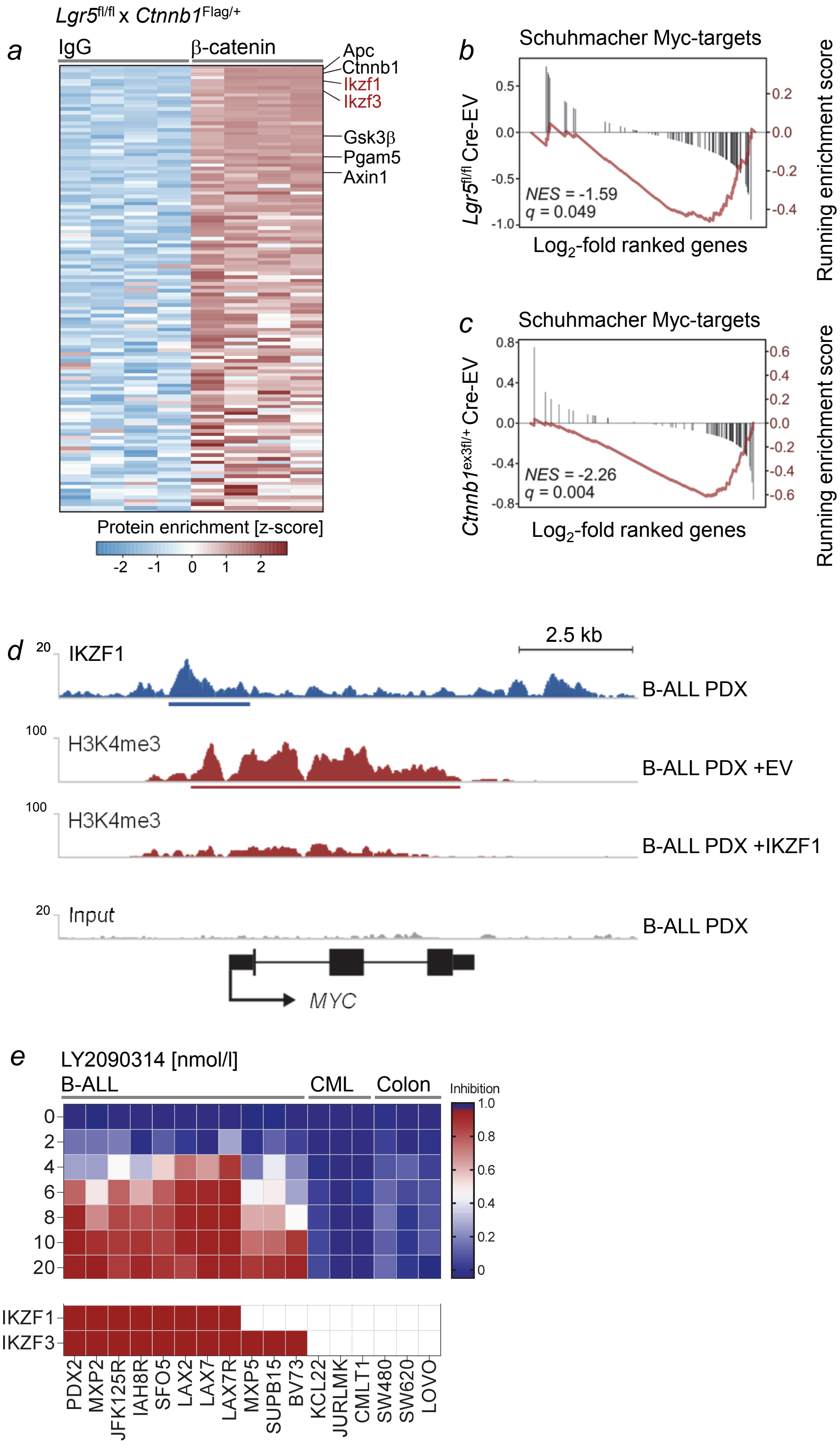


Pre-B cells from *Mb1*-Cre$^{ERT2}$ x *Lgr5*$^{fl/fl}$ x *Ctnnb1*$^{Flag/+}$ mice were transduced with *BCR-ABL1* to generate B-ALL and treated with 4-OHT for 3 days. These mice express β-catenin tagged by FLAG. Proteins bound to β-catenin were enriched by pull-down using anti β-catenin antibody. Pull-down with anti-IgG antibody was performed to normalize for non-specific binding. (**a**) Proteins bound to β-catenin and IgG antibody were analyzed by mass spectrometry and proteins significantly enriched for binding to β-catenin antibody are depicted (n=4). This analysis showed binding of β-catenin to Ikzf1 (Ikaros) and Ikzf3 (Aiolos) as top-ranking interactions (heatmap in **a**).

Gene expression changes (RNA-seq, **Fig. 3b, 3f**) in *BCR-ABL1*-transformed *Lgr5*$^{fl/fl}$ (**b**) and *Ctnnb1*$^{ex3fl/+}$(**c**) B-ALL cells upon induction of Cre-ER$^{T2}$ vs ER$^{T2}$ were studied for gene set enrichment. These gene expression changes not only showed positive enrichment of Ikzf1-targets and an Ikzf1-transcriptional program (**Fig. 6a-b**) but also negative enrichment for Myc-targets.

(**d**) Binding of IKZF1 to the *MYC* locus was studied in patient-derived B-ALL cells (LAX2) by ChIP-seq analysis (blue tracks (GSE90670). Significant binding of IKZF1 (blue) over input (gray) is underscored using ChIP-seeqer algorithm). In a different B-ALL PDX with IKZF1-deletion, ChIP-seq was performed for H3K4me3, a histone mark that denotes propensity to transcriptional activation. H3K4me3 ChIP-seq was performed in IKZF1-deleted B-ALL PDX transduced with empty vector control (EV) and B-ALL PDX after reconstitution of IKZF1.

(**e**) Responses to treatment with the GSK3β-inhibitor LY2090314 were correlated with expression of IKZF1 and IKZF3. The CML and colon cancer cell lines do not express IKZF1 and IKZF3. The B-ALL MXP2, SUPB15 and BV173 carry deletions of IKZF1 but express IKZF3.

**Extended data figure 16:** *Targeting of Lgr5⁺ B-LIC with Lgr5-ADC and GSK3β small molecule inhibitors*

*a*

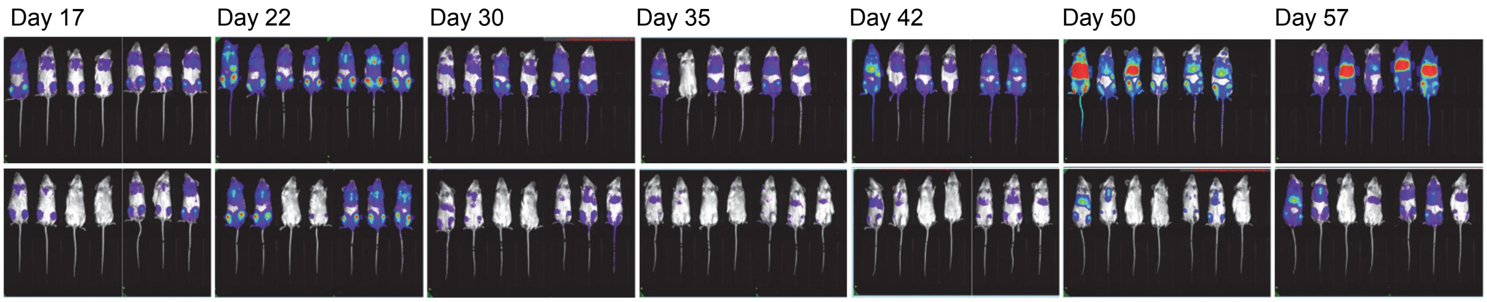


*b*

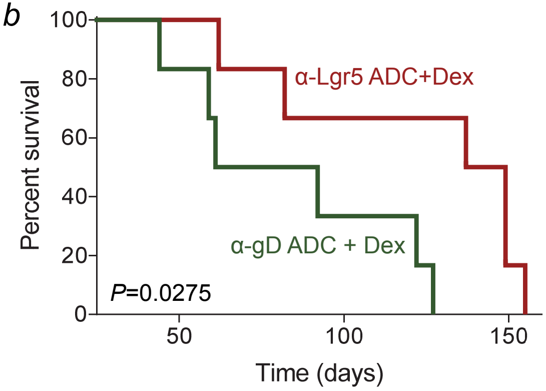


(**a**) Sub-lethally irradiated NSG mice were injected with 2 million firefly luciferase-tagged PDX2 cells (*Ph*⁺ B-ALL) and were treated with 20 mg $kg^{-1}$ anti-LGR5 ADC or control anti-gD ADC in combination with 10 mg $kg^{-1}$ dexamethasone. The leukemia progression or regression in the mice was measured by bioimaging at the indicated times. (**b**) Kaplan-Meier analysis depicts overall survival for each group (n=7) of the recipient mice (P=0.0275, log-rank test).

*c*

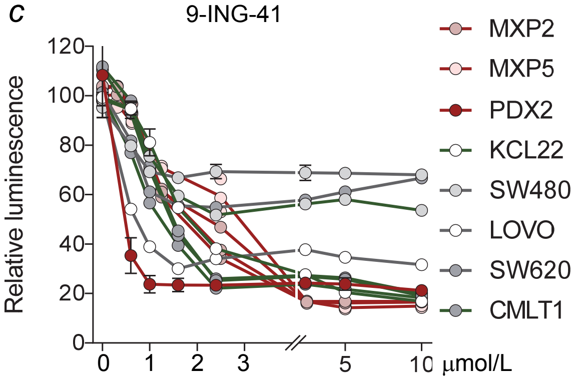


*d*

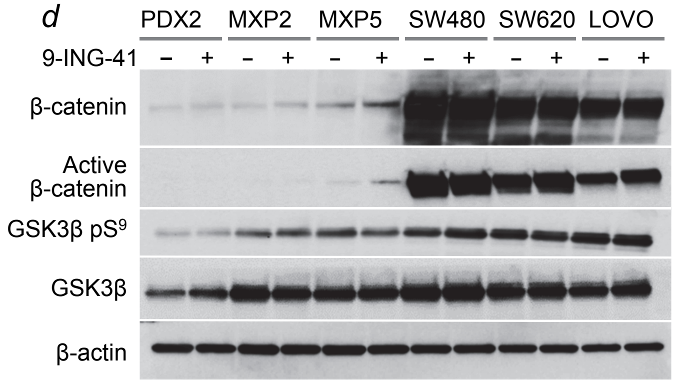


*e*

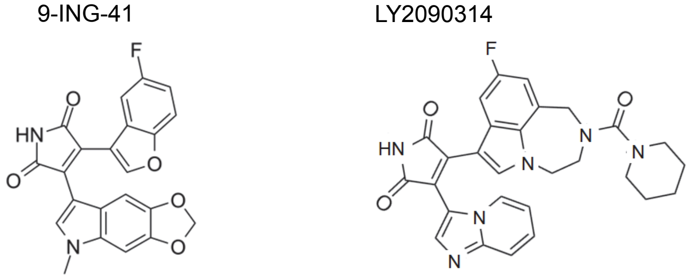


**(c)** B-ALL xenografts (red), CML (green) and colorectal cancer (grey) cell lines were treated with 0-10 $\mu mol^{-1}$ l 9-ING-41 for 3 days and relative viability was defined by measuring luminescence values.

**(d)** B-ALL xenografts and colon cancer cell lines were treated with 2 μmol $l^{-1}$ 9-ING-41 for 16 hours and Western blot analyses was performed for β-catenin, active β-catenin, GSK3β $pS^9$, GSK3β and β-actin.

**(e)** Despite structural similarities between 9-ING-41 and LY2090314, the two compounds strikingly differ with respect to their ability to induce β-catenin hyperactivation and cell death in B-ALL cells (see **Fig. 6**).
